# An inducible genetic tool for tracking and manipulating specific microglial states in development and disease

**DOI:** 10.1101/2023.12.01.569597

**Authors:** Kia M. Barclay, Nora Abduljawad, Zuolin Cheng, Min Woo Kim, Lu Zhou, Jin Yang, Justin Rustenhoven, Jose Mazzitelli Perez, Leon Smyth, Wandy Beatty, JinChao Hou, Naresha Saligrama, Marco Colonna, Guoqiang Yu, Jonathan Kipnis, Qingyun Li

**Affiliations:** Department of Neuroscience, Washington University in St. Louis School of Medicine, St. Louis, MO 63110, USA; Neuroscience Graduate Program, Washington University School of Medicine, St. Louis, MO 63110, USA; Hope Center for Neurological Disorders, Washington University School of Medicine, St. Louis, MO 63110, USA; Center for Brain Immunology and Glia (BIG), Washington University School of Medicine, St. Louis, MO 63110, USA; Department of Pathology and Immunology, Washington University in St. Louis School of Medicine, St. Louis, MO 63110, USA; Bradley Department of Electrical and Computer Engineering, Virginia Polytechnic Institute and State University, Arlington, VA 22203, USA; Immunology Graduate Program, Washington University School of Medicine, St. Louis, MO 63110, USA; Medical Scientist Training Program, Washington University School of Medicine, St. Louis, MO 63110, USA; Department of Molecular Microbiology, Washington University School of Medicine in St. Louis, School of Medicine, St. Louis, MO 63110, USA; Department of Neurology, Washington University School of Medicine in St. Louis, School of Medicine, St. Louis, MO 63110, USA; Bursky Center for Human Immunology and Immunotherapy Programs, Washington University School of Medicine in St. Louis, St. Louis, MO 63112, USA; Department of Genetics, Washington University School of Medicine in St. Louis, School of Medicine, St. Louis, MO 63110, USA

**Keywords:** Microglia, Alzheimer’s Disease, Multiple Sclerosis, Development, Single-cell RNA Sequencing, Proliferative-Region Associated Microglia, Disease-Associated Microglia, Heterogeneity, Clec7a-CreER, Plasticity, Depletion

## Abstract

Recent single-cell RNA sequencing studies have revealed distinct microglial states in development and disease. These include proliferative region-associated microglia (PAM) in developing white matter and disease-associated microglia (DAM) prevalent in various neurodegenerative conditions. PAM and DAM share a similar core gene signature and other functional properties. However, the extent of the dynamism and plasticity of these microglial states, as well as their functional significance, remains elusive, partly due to the lack of specific tools. Here, we report the generation of an inducible Cre driver line, Clec7a-CreER^T2^, designed to target PAM and DAM in the brain parenchyma. Utilizing this tool, we profile labeled cells during development and in several disease models, uncovering convergence and context-dependent differences in PAM/DAM gene expression. Through long-term tracking, we demonstrate surprising levels of plasticity in these microglial states. Lastly, we specifically depleted DAM in cuprizone-induced demyelination, revealing their roles in disease progression and recovery.

## INTRODUCTION

Microglia, the central nervous system (CNS) resident macrophages, have traditionally been characterized by their rapid immune response during injury and disease. In pathological contexts, microglia can present antigens, secrete pro- and anti-inflammatory cytokines and chemokines, and phagocytose debris and apoptotic cells.^1–3^ Recent studies, however, have explored microglia beyond their canonical immune functions. Evidence of microglia’s ability to prune synapses,^4–7^ sense and modulate neuronal activity,^8–13^ and promote myelinogenesis,^14–18^ has since implicated them as critical regulators of CNS development and homeostasis.^19–21^ Furthermore, numerous immune or microglia-specific genes have been associated with human neurological diseases, highlighting the potential disease driving roles by microglia.^22–27^ Interestingly, single-cell RNA sequencing (scRNA-seq) has revealed context-dependent microglia heterogeneity, providing new insights about the nature of their functional diversity.^28–35^ Defining the specific functions of microglial states, as well as their ability to shift between states, has been difficult due, in part, to the lack of tools to perform fate mapping and manipulate microglia in a state-specific manner.^36^

In recent years, several microglial Cre driver lines have been developed, enabling the investigation of microglial function *in vivo*. This toolbox encompasses the widely used Cx3cr1^Cre^ and Cx3cr1^CreER^ lines,^37–39^ which offer high levels of recombination efficiency, as well as the lines that are more specific to the microglial compartment such as Sall1^CreERT2^, Tmem119^CreERT2^, P2ry12^CreER^, and Cx3cr1^ccre^:Sall1^ncre^.^37,40–43^ In addition, Hexb^CreERT2^ reliably labels microglia in various disease conditions due to the stable expression of Hexb, and Cyrbb1^Cre^ demonstrates high efficiency in embryonic microglia.^44,45^ While these tools have provided valuable resources to the field, they are designed to target the microglial cell type in general and are therefore unable to drive reporter gene expression or perform cell ablation in specific microglial subpopulations.^46,47^ Given the increasing recognition of microglial heterogeneity, new genetic tools that label and manipulate distinct states of microglia with high efficiency, specificity, temporal control, and without disrupting gene function, are needed.

scRNA-seq studies have identified proliferative region-associated microglia, PAM (or axon tract-associated microglia, ATM),^28,29^ and disease-associated microglia, DAM (or microglial neurodegenerative subset, MGnD) among other states.^30,48^ PAM transiently appear in developing white matter and neurogenic niches in the early postnatal brain, and DAM are enriched in aging and neurodegenerative diseases. Spatiotemporal enrichment of these distinct states and the upregulation of certain disease risk genes, such as *Trem2* and *Apoe*, underscore their functional relevance.^28,49^ Importantly, these microglial states share a similar core gene signature, including the specific expression of C-type lectin domain containing 7A (*Clec7a*).^28,36,48^

Based on this observation, we generated an inducible Clec7a-CreER^T2^ mouse line by inserting the CreER^T2^ cassette downstream of the *Clec7a* locus without disrupting its coding region. We comprehensively characterize this model for its targeting efficiency and specificity in development, healthy adult, and mouse models of Alzheimer’s disease (AD) and multiple sclerosis (MS). Using the Clec7a-CreER^T2^ reporter line, we demonstrate its efficacy in acutely isolating PAM and DAM for scRNA-seq profiling, which reveals both convergence and context-dependent divergence of genes and pathways. Furthermore, long-term tracking of DAM following cuprizone-induced demyelination, shows that DAM are morphologically and transcriptionally plastic. Finally, through state-specific ablation in the same white matter injury model, we show that DAM are required for removing damaged myelin during demyelination in order to facilitate efficient remyelination. Collectively, we provide a versatile genetic tool and exemplify its applications to understand dynamic changes and functions of microglial states in CNS development and disease.

## RESULTS

### Generation of the Clec7a-CreER^T2^ mouse

Recent scRNA-seq studies have revealed a shared gene signature in PAM and DAM, highly enriched in the early postnatal developing white matter and neurodegenerative conditions, respectively.^28–31^ This signature, distinctive from lipopolysaccharide (LPS)-induced inflammation,^50^ involves the upregulation of a cohort of immune genes, including the anti-fungal pattern recognition receptor *Clec7a*, and the downregulation of microglial homeostatic genes (**Figure 1A**). We reasoned that *Clec7a* is an ideal marker gene to identify PAM and DAM, due to its robust and specific expression in these microglial states but not in other neuronal or glial cell types in the CNS parenchyma.^51^ To validate CLEC7A expression in PAM and DAM, we performed immunohistochemistry (IHC) on tissue sections from postnatal day 7 (P7) animals, and several disease models, namely the 5xFAD model of Alzheimer’s disease (AD), and cuprizone and experimental autoimmune encephalomyelitis (EAE) models of multiple sclerosis (MS). As expected, we observed strong and specific signals from CLEC7A antibody staining in microglial subpopulations in the P7 white matter, as well as in regions of pathology, i.e., near amyloid plaques or demyelination in the disease models (**Figure 1B and Figure S1A**).

**Figure 1.**
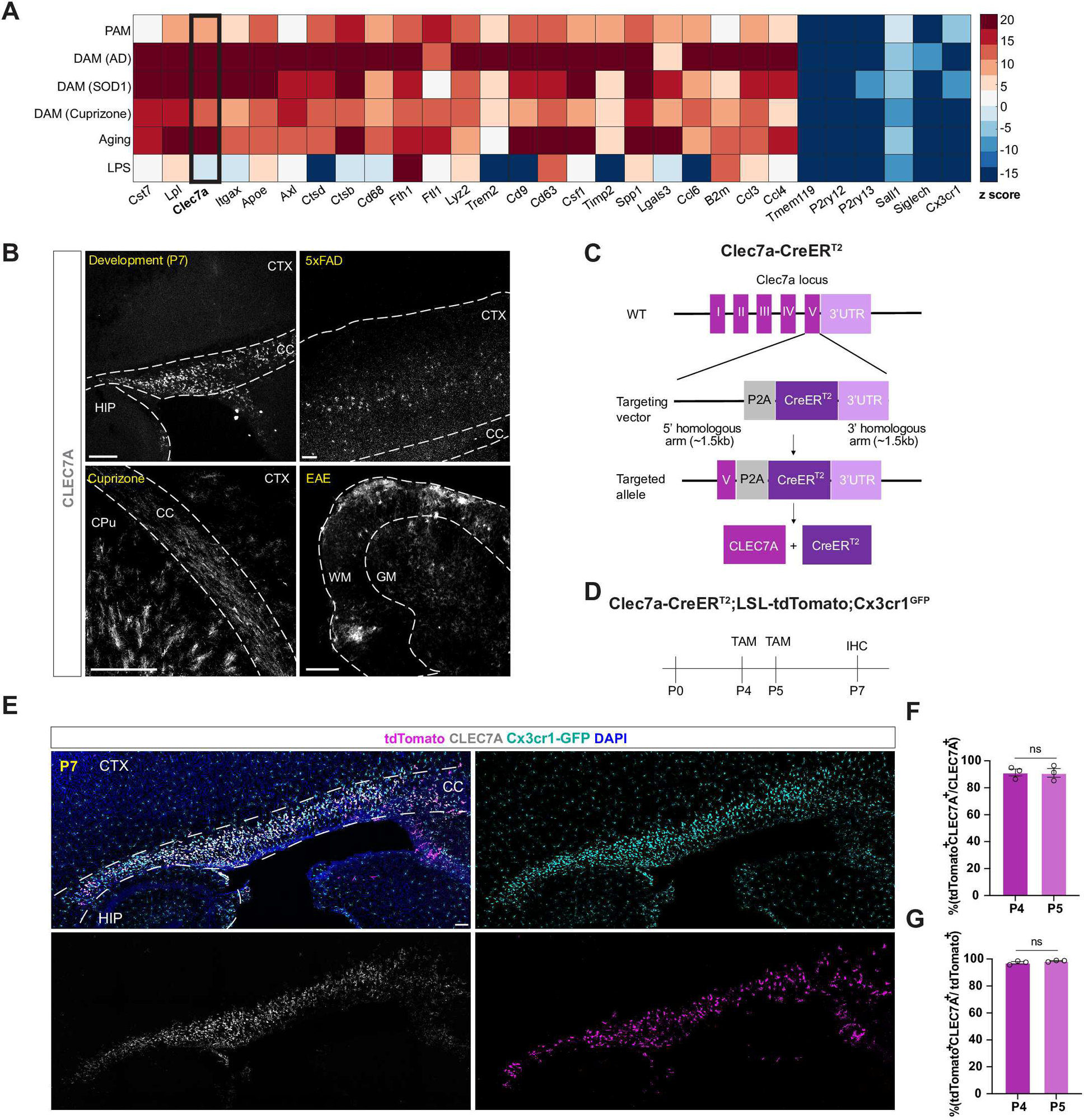
Generation of the Clec7a-CreER^T2^ mouse. **(A)** Heatmap showing the upregulation of a similar cohort of genes and downregulation of homeostatic genes in PAM during development and DAM in aging and across disease models. Input data were from previously published scRNA-seq datasets.^28–31,50^ **(B)** Immunostaining validation of CLEC7A expression in development (P7), 5xFAD, cuprizone and EAE models. Scale bar = 200um. CC, corpus callosum; CTX, cortex; HIP, hippocampus; CPu, caudate putamen; WM, white matter; GM, gray matter. **(C)** Genetic strategy for the generation of Clec7a-CreER^T2^ mouse. P2A-CreER^T2^ is inserted between exon V and 3’-UTR of *Clec7a*. The coding region remains intact. **(D)** Schematic depiction of tamoxifen (TAM) injection regimen and tissue processing timeline for PAM labeling. A single dose of TAM is injected subcutaneously on either P4 or P5, and tissues are harvested on P7. **(E)** Representative IHC images showing efficient and specific labeling of PAM in the corpus callosum of a P7 brain in the Clec7a-CreER^T2^;LSL-tdTomato;Cx3cr1^GFP^ mouse (triple heterozygous). tdTomato and CLEC7A signals largely overlap, and these cells are also GFP^+^. Scale bar = 100um. CC, corpus callosum; CTX, cortex; HIP, hippocampus. **(F)** Quantification of labeling efficiency (percentage of tdTomato^+^CLEC7A^+^/CLEC7A^+^ microglia) of PAM in P7 corpus callosum following tamoxifen injection at P4 or P5, n = 3 animals per group (3 sections per animal). Student’s T-test, ns = not significant. Error bars represent mean +/− SEM. **(G)** Quantification of labeling specificity (percentage of tdTomato^+^CLEC7A^+^/tdTomato^+^ microglia) of PAM in P7 corpus callosum following tamoxifen injection at P4 or P5, n = 3 animals per group (3 sections per animal). Student’s T-test, ns = not significant. Error bars represent mean +/− SEM. **See also Figure S1.**

This prompted us to create a new Cre driver mouse line utilizing the *Clec7a* regulatory regions to specifically target PAM and DAM. Through CRISPR-Cas9 genome editing, we inserted the P2A-CreER^T2^ cassette between the last exon and 3’-UTR of the *Clec7a* gene locus (Clec7a-CreER^T2^). This resulted in an inducible Cre under the control of the endogenous regulatory elements of *Clec7a* without disrupting its coding region (**Figure 1C**). Southern blots confirmed no other random insertions (**Figure S1B**). Upon tamoxifen administration, this strain is expected to induce Cre-mediated recombination leading to the expression of reporters, conditional alleles or cell ablation systems in PAM and DAM.

To test whether Clec7a-CreER^T2^ can drive reporter expression in PAM, we crossed it with LSL-tdTomato (Ai14) and Cx3cr1^GFP^ mice. We administered tamoxifen to the pups on either P4 or P5 when PAM start to appear in the developing white matter regions^28,29^ (**Figure 1D**). We indeed observed tdTomato^+^ cells in the corpus callosum and cerebellar white matter at P7 when PAM are at their peak density (**Figure 1E and Figure S1C**). Quantification of labeling efficiency demonstrated that up to ∼90% of CLEC7A^+^(also GFP^+^) microglia were co-labeled with tdTomato (**Figure 1F**). tdTomato^+^ cells were almost all CLEC7A^+^GFP^+^ (**Figure 1G**). We found no spontaneous labeling in control mice (Clec7a-CreER^T2^;LSL-tdTomato) without tamoxifen injection (**Figure S1D**).

Because PAM (and DAM) also upregulate *Itgax* (CD11c),^18,28,30^ we performed a similar experiment using the existing CD11c-CreERT line as a comparison. Although this model was able to label PAM in the corpus callosum, it drove a more widespread reporter expression pattern in the P7 microglia, including labeling CD11c^−^ cells in the ventral striatum, suggesting decreased specificity (**Figures S1E, S1F**). Taken together, these data suggest that our newly generated Clec7a-CreER^T2^ line targets PAM at high efficiency and specificity.

### Clec7a-CreER^T2^ does not affect *Clec7a* gene expression or function

As such genomic manipulations may cause haploinsufficiency of a gene and *Clec7a* has been shown to play critical roles in certain disease contexts,^45,47,52,53^ we wanted to examine whether CreER^T2^ insertion affects *Clec7a* gene expression and function. We found no significant differences in the level of CLEC7A protein in PAM between homozygous, heterozygous and wildtype littermates of the transgenic mice (**Figures 2A, 2B**). The number, morphology, or regional distribution of PAM were also comparable regardless of the genotypes (**Figures 2C, 2D and Figure S2**).

**Figure 2.**
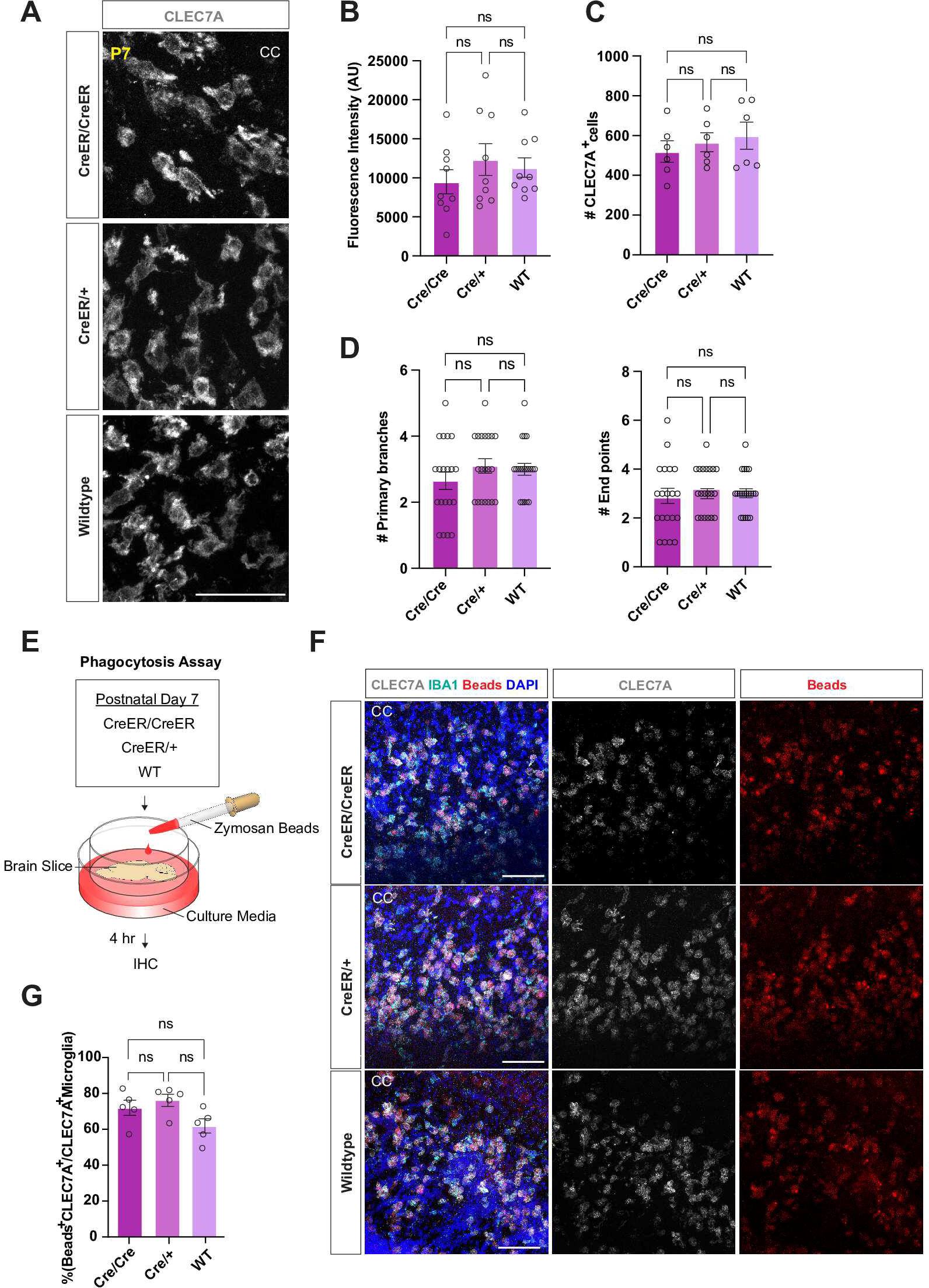
Clec7a-CreER^T2^ does not affect *Clec7a* gene expression or function. **(A)** Immunostaining of CLEC7A in the corpus callosum of Clec7a-CreER^T2^ homozygous, heterozygous and wildtype P7 littermates. Scale bar = 50um. **(B)** Quantification of CLEC7A fluorescence intensity across Cre genotypes. n = 9 sections from 3 mice. One-way ANOVA with Tukey’s multiple comparison test. ns: not significant. Error bars represent mean +/− SEM. **(C)** Quantification of the number of CLEC7A^+^ cells across Cre genotypes. n = 5-6 mice (3 sections per animal). One-way ANOVA with Tukey’s multiple comparison test. ns: not significant. Error bars represent mean +/− SEM. **(D)** Quantification of the numbers of primary branches and end points per microglia across Cre genotypes. n =20. One-way ANOVA with Tukey’s multiple comparison test. ns: not significant. Error bars represent mean +/− SEM. **(E)** Schematic depiction of the experimental design for phagocytosis assay. **(F)** Immunostaining of CLEC7A^+^ microglia (PAM) overlapping with engulfed pHrodo zymosan beads across Cre genotypes. Scale bar = 100 um; CC, corpus callosum. **(G)** Quantification of the percentage of CLEC7A^+^ microglia (PAM) phagocytosing pHrodo zymosan beads in corpus callosum (CC) across Cre genotypes. n = 5. One-way ANOVA with Tukey’s multiple comparison test. ns: not significant. Error bars represent mean +/− SEM. **See also Figure S2.**

To determine whether Cre insertion affects *Clec7a* function, acute brain slices of Cre homozygous, heterozygous and wildtype P7 littermates were incubated with pH-sensitive beads coated with zymosan, which is a ligand of CLEC7A.^54^ Phagocytosis of the beads was detected by IHC (**Figure 2E**). We observed no differences in the numbers of beads engulfed by PAM between genotypes (**Figures 2F, 2G**). Therefore, we concluded that the Clec7a-CreER^T2^ mouse has intact *Clec7a* gene function and is suitable for studying *Clec7a^+^* microglial states, such as PAM.

### Clec7a-CreER^T2^ targets CNS border and peripheral myeloid cells to a lesser extent

To further characterize the immune populations targeted by Clec7a-CreER^T2^, we performed high-dimensional flow cytometry on P7 brain parenchyma and border regions, including dura, leptomeninges, and choroid plexus in the reporter mice (**Figure 3A and Figure S3A**). We quantified tdTomato labeling of a range of immune cells within each tissue compartment, including both myeloid and lymphoid populations. Consistent with the histology data, tdTomato labeling was restricted to microglia in the parenchymal tissue, with up to ∼1,000 cells recovered per P7 sample (**Figure 3B**). Smaller groups of tdTomato^+^ cells were observed in the choroid plexus and meninges, consisting primarily of macrophages and other myeloid cells (**Figure 3B**). Immunohistochemistry on flat-mounted choroid plexus, leptomeninges, and dura confirmed the presence of tdTomato^+^CD206^+^ macrophages in these tissues (**Figures 3C-E**). Flow cytometry analysis of adult tissues showed a similar pattern of labeling in the choroid plexus and meninges, with very few labeled microglia in the brain parenchyma as expected (**Figures S3C-E**). Peripheral myeloid labeling was also observed in the blood and liver (**Figures S3B and S3E**). These data suggest that the Clec7a-CreER^T2^ model predominantly targets microglial subpopulations in the healthy brain, and to a lesser extent CNS border-associated myeloid cells.

**Figure 3.**
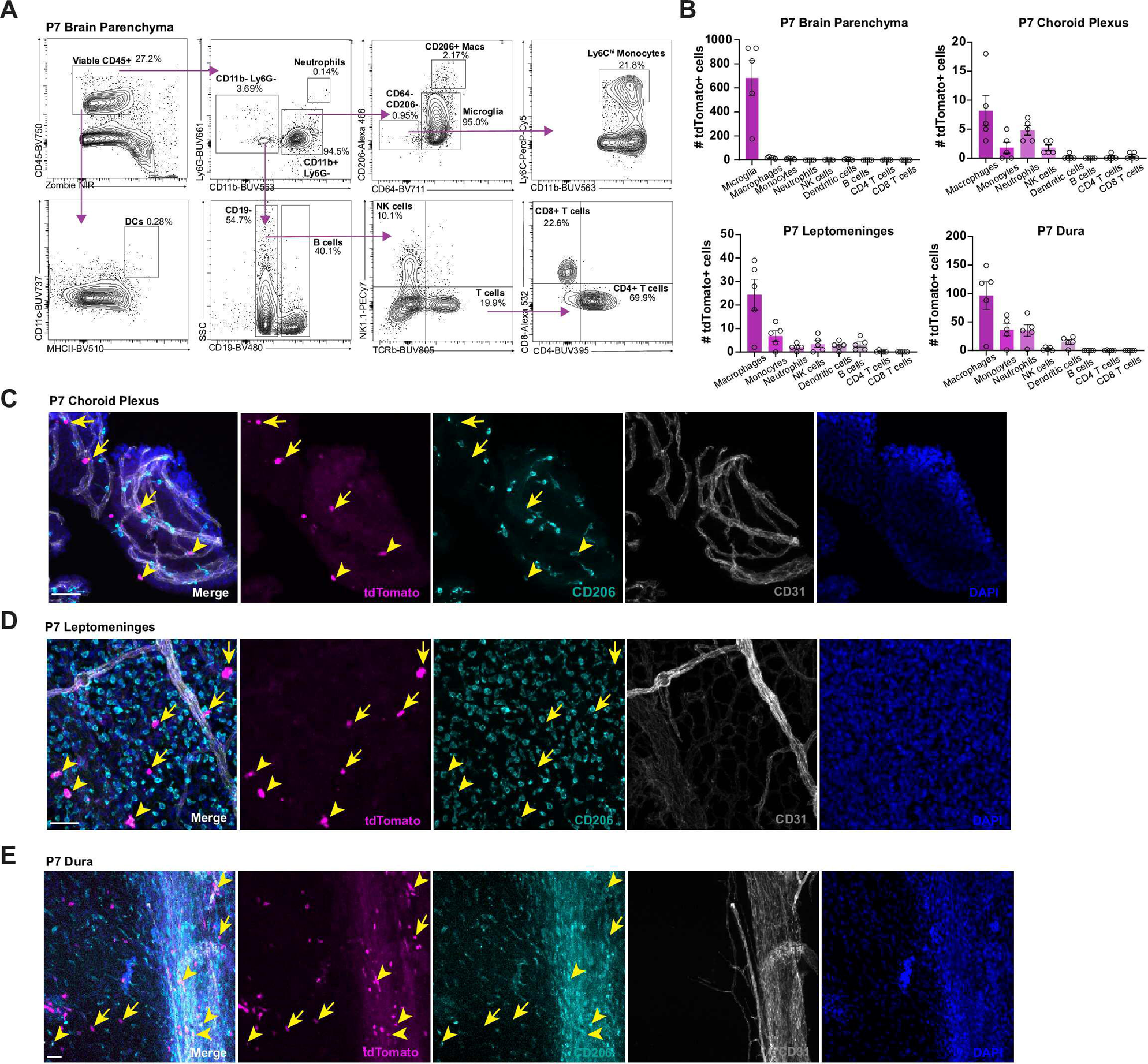
Clec7a-CreER^T2^ targets CNS border and peripheral myeloid cells to a lesser extent. **(A)** Flow cytometry gating strategy for different immune cells in the Clec7a-CreER^T2^;LSL-tdTomato mice (represented by P7 brain parenchymal tissue). **(B)** Quantification of tdTomato^+^ immune cells in P7 brain parenchyma and borders. n =5 mice. Error bars represent mean +/− SEM. **(C)** Validation of tdTomato^+^ macrophages and immune cells in P7 choroid plexus. Arrowheads represent tdTomato^+^CD206^+^ (macrophages) and arrows represent tdTomato^+^CD206^−^ immune cells. Scale bar = 50um. **(D)** Validation of tdTomato^+^ macrophages and immune cells in P7 leptomeninges. Arrowheads represent tdTomato^+^CD206^+^ (macrophages) and arrows represent tdTomato^+^CD206^−^ immune cells. Scale bar = 50um. **(E)** Validation of tdTomato^+^ macrophages and immune cells in P7 dura mater. Arrowheads represent tdTomato^+^CD206^+^ (macrophages) and arrows represent tdTomato^+^CD206^−^ immune cells. The large vessel is the sinus. Scale bar = 50um. **See also Figure S3.**

### Clec7a-CreER^T2^ reporter labels DAM across disease models

Given that *Clec7a* is a shared marker gene in neurodegenerative disease conditions, we next asked whether Clec7a-CreER^T2^ reporter mice could be used to label disease-associated microglia. We turned to three different disease models: 5xFAD, cuprizone-induced demyelination, and EAE (**Figure 4A**).

**Figure 4.**
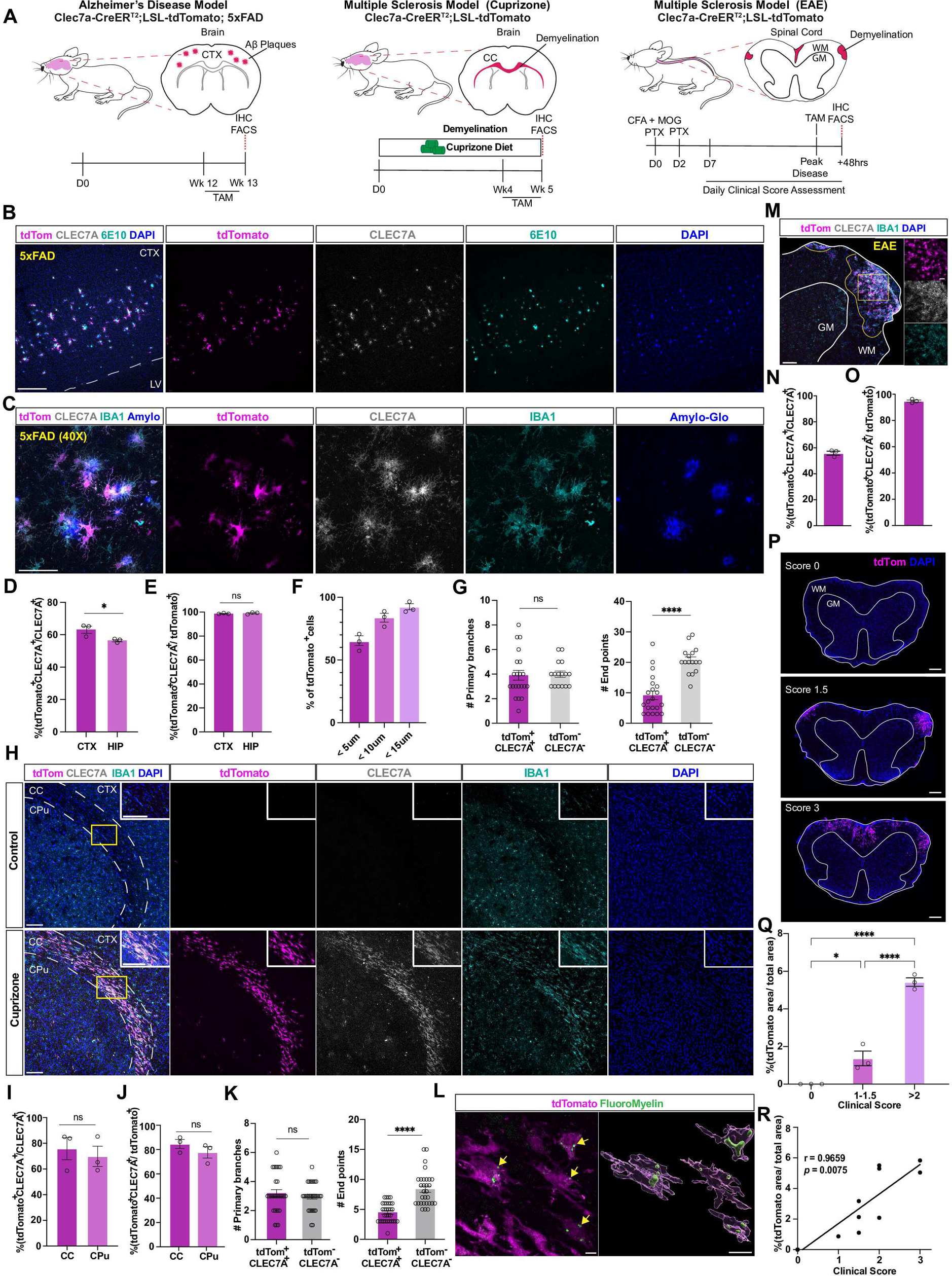
Clec7a-CreER^T2^ reporter labels DAM across disease models. **(A)** Schematic of three disease models, 5XFAD, cuprizone and EAE, with corresponding experimental design. **(B)** Representative image showing tdTomato^+^CLEC7A^+^ microglia surrounding amyloid plaques (6E10^+^) in Clec7a-CreER^T2^;LSL-tdTomato;5xFAD mice. Scale bar = 250um. **(C)** Higher magnification image showing tdTomato^+^CLEC7A^+^ microglia clustering around amyloid plaques (Amylo-glo^+^) in the cortex of Clec7a-CreER^T2^;LSL-tdTomato;5xFAD mice. Scale bar = 50 um. **(D)** Quantification of labeling efficiency (percentage of tdTomato^+^CLEC7A^+^/CLEC7A^+^ microglia) following tamoxifen administration in the 5xFAD model. **(E)** Quantification of labeling specificity (percentage of tdTomato^+^CLEC7A^+^/tdTomato^+^ microglia) following tamoxifen administration in the 5xFAD model. **(F)** Percentage of tdTomato^+^ microglia found within 5-, 10- and 15-micron distance from amyloid plaques. **(G)** Quantification of the numbers of primary branches (left) and end points (right) in the microglia from the 5xFAD Cre reporter mice. n = 15-20 per group. **(H)** Representative image showing tdTomato^+^CLEC7A^+^ microglia in control (top) and cuprizone (bottom) treated mice. Scale bar = 100um. **(I)** Quantification of labeling efficiency (percentage of tdTomato^+^CLEC7A^+^/CLEC7A^+^ microglia) following tamoxifen administration in the cuprizone model. **(J)** Quantification of labeling specificity (percentage of tdTomato^+^CLEC7A^+^/tdTomato^+^ microglia) following tamoxifen administration in the cuprizone model. **(K)** Quantification of the numbers of primary branches (left) and end points (right) in the microglia from the cuprizone Cre reporter mice. n = 25-30 per group. **(L)** Representative image and 3D reconstruction of tdTomato^+^ cells with intracellular myelin fragments (FluoroMyelin^+^) indicated by yellow arrows. Scale bars = 5um (left) and 10um (right). **(M)** Representative image showing tdTomato^+^CLEC7A^+^ microglia in the spinal cord of EAE Cre reporter mice. Scale bar = 100um and 25um (zoomed-in). **(N)** Quantification of labeling efficiency (percentage of tdTomato^+^CLEC7A^+^/CLEC7A^+^ microglia) following tamoxifen administration in the EAE model. **(O)** Quantification of labeling specificity (percentage of tdTomato^+^CLEC7A^+^/tdToamto^+^ microglia) following tamoxifen administration in the EAE model. **(P)** Representative images of tdTomato labeling in the spinal cord of EAE mice with clinical scores of 0 (top), 1.5 (middle) and 3 (bottom). Scale bar = 1000um. **(Q)** Quantification of the percentage of tdTomato^+^ area in the spinal cord across clinical scores. n = 3 mice per group. **(R)** Pearson correlation between the percentage of tdTomato^+^ area in spinal cord and clinical scores. Each data point represents an animal. For (D-F) and (I-K; N-O), n = 3 mice per group (3 sections per animal). Quantifications performed for labeling efficiency, specificity and morphology were calculated by Student’s T-test. For (Q and R), quantifications performed for EAE labeling in relation to clinical score were calculated by one-way ANOVA with Tukey’s multiple comparison test. ns = not significant, * *p* < 0.05, ** *p* < 0.01, *** *p* < 0.001, **** *p* < 0.0001. Error bars represent mean +/− SEM. CTX, cortex; LV, lateral ventricle; WM, white matter; GM, gray matter; CC, corpus callosum; CPu, caudate putamen. **See also Figure S4.**

First, we generated Clec7a-CreER^T2^;LSL-tdTomato;5xFAD mice to label DAM in a model of amyloidosis. Tamoxifen injections at 3 months of age labeled up to ∼60% of CLEC7A^+^ microglia in the cortex and hippocampus, sites of amyloid accumulation in these mice (**Figures 4B-D**). Among labeled cells, 98-99% were CLEC7A^+^, indicating high labeling specificity (**Figure 4E**). Labeled microglia appeared in the vicinity of 6E10^+^ and Amylo-Glo^+^ plaques as expected for DAM, with 64.3% located fewer than 5um away from a plaque and 91.8% appearing fewer than 15um away (**Figure 4F**). In addition, labeled microglia were less ramified than tdTomato^−^CLEC7A^−^ microglia, with fewer terminal branches (**Figure 4G**), consistent with previous descriptions of DAM in a reactive state. Again, no spontaneous recombination was seen even in 6-month-old animals (**Figure S4A**). These data suggest that Clec7a-CreER^T2^ mice specifically label DAM in a model of Alzheimer’s disease.

Next, we tested whether our reporter mouse can label DAM in two white matter disease models. In the first model, adult mice were fed a diet containing 0.2% cuprizone for 5 weeks to induce acute demyelination in heavily myelinated regions (**Figure S4B**).^55^ Tamoxifen injected during the last week of treatment (**Figure 4A**) labeled ∼70% of CLEC7A^+^ microglia in the corpus callosum (**Figures 4H, 4I**) and white matter tracts within the caudate putamen (**Figure S4C**). Notably, these labeled cells were also positive for other genes known to be upregulated in DAM, such as CD68 and CD11c (**Figures S4D, S4E**). Labeling in the cortex was minimal as expected (**Figure S4F**). The labeling specificity was close to 80% (**Figure 4J**), a bit lower than that in the developmental or AD conditions, presumably due to spontaneous remyelination in this model. Similar to the DAM morphology seen in the AD model, tdTomato^+^CLEC7A^+^ microglia displayed fewer branches in the cuprizone model (**Figure 4K**). Consistent with a functionally reactive state, these labeled cells often contained myelin inclusions, suggesting phagocytosis of myelin debris (**Figure 4L**).

To further explore the versatility of the Clec7a-CreER^T2^ reporter mouse, we used the classical MS model, reactive EAE, which manifests its pathology in the spinal cord. To induce EAE, adult Clec7a-CreER^T2^;LSL-tdTomato mice were immunized with an emulsion of complete Freund’s adjuvant (CFA) and myelin oligodendrocyte glycoprotein (MOG), followed by injections of pertussis toxin (PTX) (**Figure 4A**). Beginning one week after immunization, animals were assessed daily for signs of motor deficits. At peak disease score, animals were injected with one dose of tamoxifen and sacrificed 48 hours later for histology analysis. On average, the labeling efficiency was close to 60% with over 90% specificity (**Figures 4M-O**). Interestingly, the percentage of tdTomato labeled cells positively correlated with the disease scores (**Figures 4P-R**), suggesting that the extent of labeling can be used as a readout of pathology. It is worth mentioning that certain peripheral myeloid cells would also be labeled (**Figures S3C-E**), and therefore tdTomato^+^ cells in this model included both microglia and infiltrated myeloid cells that express *Clec7a*.

Collectively, these results demonstrate that our Clec7a-CreER^T2^ reporter line targets DAM-like states in a variety of disease models both in the brain and spinal cord.

### Isolation of microglial subpopulations by the reporter line for scRNA-seq

Despite the consensus that PAM and DAM upregulate many common genes (**Figure 1A**), systematic comparisons between these similar microglial states have been challenging due to the rarity of these cells in each condition and different isolation protocols used to generate the existing datasets.^56^ We reasoned that we can employ the Clec7a-CreER^T2^ reporter mice to acutely isolate various microglial states and perform scRNA-seq to define the convergent and divergent genes and pathways underlying each biological context. We used a previously published microglia isolation protocol^57^ and fluorescence activated cell sorting (FACS) to collect tdTomato^+^ and tdTomato^−^ microglia, from P7 and 5xFAD brains as well as EAE spinal cords (**Figures 5A and 5B**). Plate-based deep scRNA-seq and high dimensional clustering analysis identified 10 distinct microglial clusters: early postnatal homeostatic (C0), adult brain homeostatic (C1), adult spinal cord homeostatic (C2), PAM (C3), transitional DAM (C4) which clustered between homeostatic microglia and DAM clusters, DAM1 (C5) which expressed DAM markers but at lower levels, DAM2 (C6), MHCII^+^ (C7), IEG^+^ (C8) and Dividing cluster (C9) (**Figures 5C, 5F, 5G and Table S1**).

**Figure 5.**
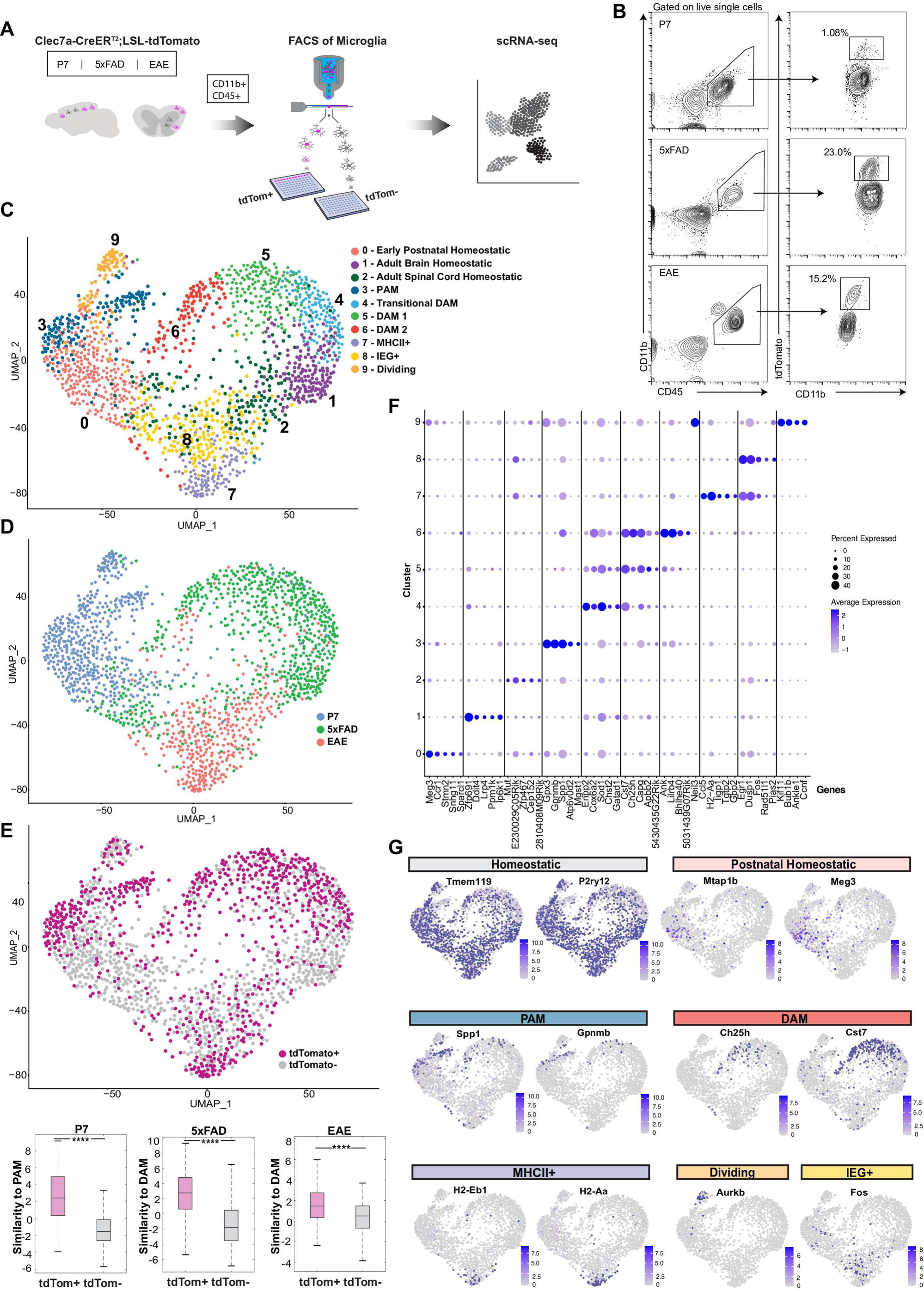
scRNA-seq analysis of PAM and DAM. **(A)** Schematic of experimental design for microglial isolation using the reporter mice to do scRNA-seq across developmental and disease conditions. **(B)** Flow cytometry gating for isolating tdTomato^+^ microglia by experimental conditions. **(C)** UMAP plot showing 10 microglia clusters from tdTomato^+^ and tdTomato^−^ cells in P7, 5xFAD and EAE mice. IEG: immediate early genes. **(D)** UMAP plot showing microglia clusters split by condition. **(E)** UMAP plot showing microglia clusters split by tdTomato expression (top) and box plots showing the similarity of tdTomato^+^ and tdTomato^−^ cells to PAM or DAM in different conditions (bottom) based on signature gene enrichment, **** *p* < 0.0001. **(F)** Dot plots showing representative marker gene expression for each microglia cluster in Figure 5C. **(G)** Features plots highlighting representative marker genes to distinguish selective microglial states. **See also Table S1.**

These clusters formed different domains on UMAP according to experimental conditions (**Figure 5D**). Interestingly, each condition contained its own homeostatic microglial cluster (*Tmem119^+^*, *P2ry12^+^*), i.e. cluster 0 for P7 (also *Mtap1b^+^*, *Meg3^+^*), cluster 1 for 5xFAD and cluster 2 for EAE, reflecting the differences of the microenvironments from the developing brain, adult brain and adult spinal cord, respectively (**Figures 5C-D and 5F-G**). Each condition also had a predominant context-dependent reactive state: PAM (cluster 3) for P7, DAM (cluster 4, 5, 6) for 5xFAD, and MHCII^+^ (cluster 7) for EAE, which were marked by distinct signature genes (**Figures 5D, 5F and 5G**).

Importantly, tdTomato^+^ and tdTomato^−^ cells were also segregated on UMAP (**Figure 5E**). Furthermore, tdTomato^+^ cells from P7 were more similar to the previously characterized PAM state than P7 tdTomato^−^ cells, whereas tdTomato^+^ cells from EAE and 5xFAD, had higher levels of similarity to DAM than tdTomato^−^ cells from each condition (**Figure 5E**). These data suggest effective enrichments of *Clec7a*^+^ reactive microglial states by the reporter line.

### scRNA-seq reveals both convergent and divergent genes and pathways in reactive microglial states

We decided to focus our analysis on PAM and DAM-related clusters. Based on the published gene expression datasets for PAM and DAM,^28,30^ cluster 3 showed the highest similarity score (METHODS) to PAM. This was followed by DAM-related clusters (cluster 6, 5 and 4) and to a lesser extent homeostatic clusters. On the other hand, cluster 6 was the most similar to the previously described DAM state, followed by PAM (cluster 3), and other DAM-related clusters (**Figures 6A-B**).

**Figure 6.**
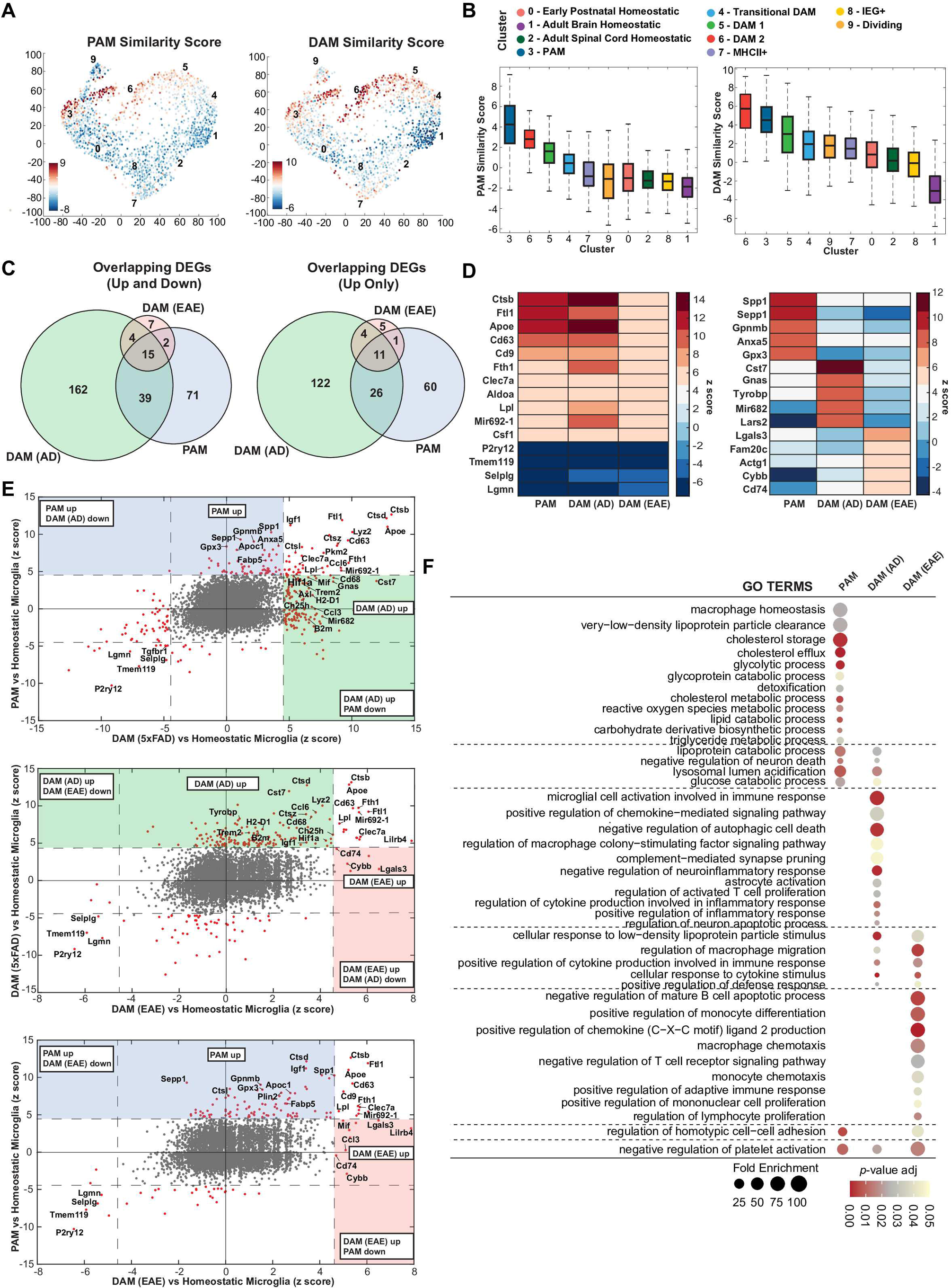
scRNA-seq reveals convergent and distinct gene signatures between PAM and DAM. **(A)** Heatmap representing similarity of each cluster to PAM (left) and DAM (right) projected on microglia UMAP plot. **(B)** Box plots showing similarity to PAM (left) and DAM (right) by cluster. **(C)** Venn diagram showing the numbers of unique and overlapping DEGs for PAM(P7), DAM(5xFAD) and DAM(EAE) compared to condition-specific homeostatic microglia. Left panel includes both up- and down-regulated genes and right panel only includes upregulated genes. **(D)** Heatmap showing common upregulated and downregulated genes (left) and condition-specific upregulated genes (right) in PAM(P7), DAM(5xFAD) and DAM(EAE) microglia. **(E)** Gene expression changes (relative to homeostatic microglia) in one condition compared to another; PAM(P7) vs DAM(5xFAD) – top, DAM (5xFAD) vs DAM (EAE) – middle, PAM(P7) vs DAM (EAE) – bottom. Genes showing significant change are in red. **(F)** Shared and unique GO terms identified for each condition. Fold enrichment represented by circle size and adjusted *p*-value represented by color scale. **See also Tables S2-4.**

To define the shared core signature and differences between these similar microglial states, we subset cells from the clusters that showed the highest similarities scores to PAM (cluster 3) and DAM (cluster 6). For each condition, we identified differentially expressed genes (DEGs) between the context-specific reactive state and homeostatic state. We found 15 shared DEGs, with 11 upregulated genes (*Clec7a*, *Lpl*, *Cd63*, *Cd9*, *Apoe*, *Csf1*, *Ctsb*, *Ftl1*, *Fth1*, *Aldoa*, *Mir692-1*) and 4 downregulated genes (*P2ry12*, *Tmem119*, *Selplg*, *Lgmn*) in all 3 conditions (**Figures 6C, 6D and Table S2**). Interestingly, each condition also displayed unique signatures, such as *Spp1*, *Gpnmb*, *Gpx3* for P7, *Cst7*, *Tyrobp, Mir682* for 5xFAD, and *Cd74*, *Lgals3*, *Cybb* for EAE (**Figures 6C, 6D**). Pairwise comparisons provided a comprehensive view of DEGs that were shared or associated with one condition but not the other (**Figure 6E and Table S3**).

To determine how upregulated genes may inform convergent and divergent functional processes in a given condition, we performed GO term analysis. Many upregulated genes in P7 were relevant for metabolic processes including cholesterol and lipid metabolism. Genes upregulated in 5xFAD were primarily involved in microglial cell activation and immune response. Interestingly, complement-mediated synapse pruning, astrocyte activation and regulation of activated T cell proliferation were among the enriched terms as well. EAE genes were involved in regulation of adaptive immune response, lymphocyte proliferation and monocyte differentiation/chemotaxis. GO terms shared between P7 and 5xFAD included negative regulation of neuron death, and lysosomal lumen acidification. GO terms shared between P7 and EAE were limited to regulation of homotypic cell-cell adhesion. GO terms shared between 5xFAD and EAE included regulation of macrophage migration and positive regulation of cytokine production involved in immune response. Interestingly, all three conditions shared the term, negative regulation of platelet activation (**Figure 6F and Table S4).** These data suggest shared and distinct functions between PAM and DAM states, which may underlie their roles in development and disease.

### Long-term tracking of DAM reveals microglial state plasticity

Once in a reactive state such as DAM, the long-term fate of these microglia remains an outstanding question in the field.^36^ To address this, we used our Clec7a-CreER^T2^;LSL-tdTomato reporter mouse to track DAM in a white matter disease model where pathology can be resolved. We leveraged the cuprizone model since cessation of cuprizone diet allows for oligodendrocyte precursor cell proliferation, differentiation and remyelination. Adult mice were fed cuprizone to induce demyelination and injected with tamoxifen to label DAM as shown earlier. After five weeks of treatment, animals were switched to a control diet for two additional weeks to allow for spontaneous remyelination (**Figure 7A**). Histology analysis showed that tdTomato^+^ microglia labeled during demyelination were still present in the white matter tracts after two weeks of remyelination (**Figure 7B**). Interestingly, these cells were mostly negative for CLEC7A (**Figures 7B, 7C**). These data suggest that microglia, once activated, may survive long term in the brain and downregulate disease-associated genes such as *Clec7a* during resolution.

**Figure 7.**
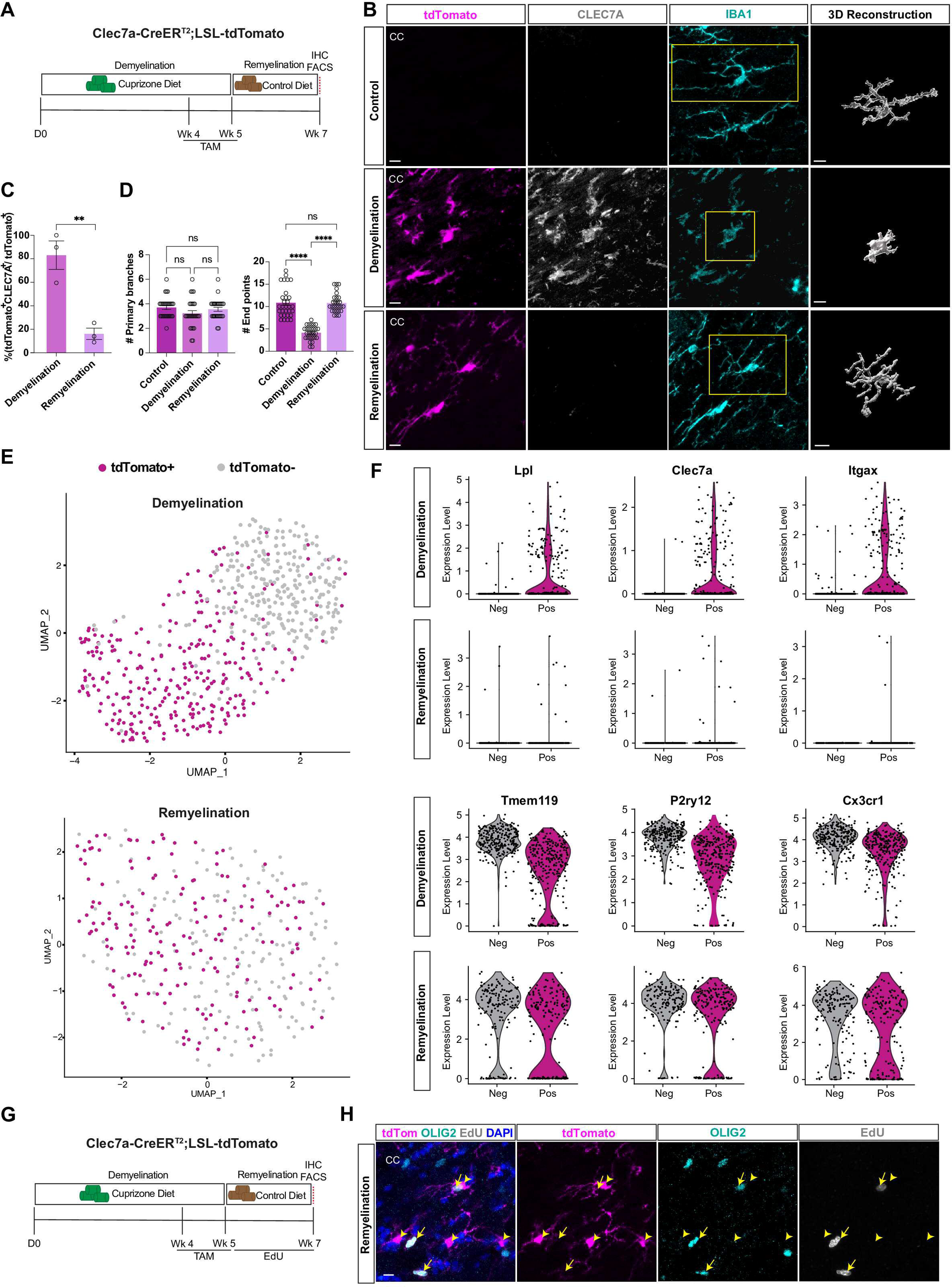
DAM return to homeostasis during remyelination in cuprizone model. **(A)** Schematic of experimental timeline for fate mapping of DAM in cuprizone model. **(B)** Representative immunostaining images showing microglia labeled by the Cre reporter, CLEC7A and IBA1 in control (top), demyelination (middle) and remyelination (bottom) conditions. 3D reconstruction of boxed microglia (right most column) showing morphological changes. Scale bar = 10um. CC, corpus callosum. **(C)** Quantification for the percentage of CLEC7A^+^ microglia among tdTomato^+^ microglia in demyelination and remyelination. n = 3 mice per group (3 sections per animal). Student’s T-test. ** *p* < 0.01. Error bars represent mean +/− SEM. **(D)** Quantification for the numbers of primary branches (left) and end points (right) in control (tdTomato^−^CLEC7A^−^), demyelination (tdTomato^+^CLEC7A^+^) and remyelination (tdTomato^+^CLEC7A^−^) conditions. n = 25-30. One-way ANOVA with Tukey’s multiple comparison test. ns: not significant, **** *p* < 0.0001. Error bars represent mean +/− SEM. **(E)** UMAP plot showing tdTomato^+^ and tdTomato^−^ microglia separated into distinct clusters during demyelination (top), and intermingled following remyelination (bottom). **(F)** Violin plots showing expression of DAM (*Lpl*, *Clec7a*, *Itgax*) and homeostatic (*Tmem119*, *P2ry12*, *Cx3cr1*) marker genes split by tdTomato expression after demyelination and remyelination. **(G)** Schematic of experimental design for EdU labeling during remyelination. **(H)** Representative images showing tdTomato^+^ microglia and OLIG2^+^ cells with EdU labeling during remyelination. Arrows indicate OLIG2^+^EdU^+^ cells and arrowheads indicate tdTomato^+^ microglia negative for EdU. Scale bar = 50um. **See also Figure S5.**

To further determine if the DAM phenotype was fully reverted, we sorted and sequenced tdTomato^+^ and tdTomato^−^ cells at the demyelination (5-week cuprizone diet) and remyelination (2-week control diet) stages. Again, tdTomato^+^ cells isolated during demyelination were in accordance with the DAM state upregulating marker genes such as *Lpl*, *Clec7a* and *Itgax*, compared to tdTomato^−^ homeostatic microglia (**Figures 7E, 7F**). In contrast, tdTomato^+^ and tdTomato^−^ cells isolated during remyelination were no longer separated into distinct clusters (**Figure 7E**). tdTomato^+^ microglia downregulated DAM marker genes and upregulated homeostatic genes including *Tmem119*, *P2ry12* and *Cx3cr1* to similar levels as tdTomato^−^ microglia (**Figure 7F and Figure S5A**). Concomitant with these gene expression changes, the morphology of tdTomato^+^ microglia also shifted from the amoeboid morphology during demyelination to a ramified, homeostatic morphology during remyelination (**Figures 7B, 7D**). These data suggest that DAM in the cuprizone model return to the homeostatic state after remyelination.

Because this fate mapping strategy not only traces the microglia initially activated upon demyelination injury but also their progenies, we asked whether it is the same DAM population or the daughter cells that undergoes this phenotypic conversion. To address this question, we injected EdU to the fate mapping model throughout the remyelination phase to mark any potential microglial proliferation (**Figure 7G**). Strikingly, we found no overlap between tdTomato^+^ microglia and EdU (**Figure 7H and Figure S5B**), suggesting minimal levels of cell proliferation in the labeled population during remyelination. This is consistent with microgliosis predominantly occurring with disease pathology. In contrast, we observed abundant OLIG2^+^ EdU^+^ cells due to an increase in oligodendrocyte lineage proliferation for myelin repair (**Figure 7H**). Therefore, it is the exact same microglial cells that are converted to homeostasis during resolution.

Taken together, these data suggest the Clec7a-CreER^T2^ driver line is suitable for long term tracking of specific microglial states, through which we demonstrate that microglia can alternate between the reactive and homeostatic states, both transcriptionally and morphologically, depending on the context.

### Microglial state-specific depletion demonstrates protective roles of DAM for the recovery of cuprizone-induced demyelination

Microglial depletion through pharmacological or genetic agents remains one of the most effective approaches to study microglial function in a biological process^58–60^. These studies generally either target the microglial survival factor, Colony Stimulating Factor 1 Receptor (CSF1R), or use pan-microglial drivers to drive a suicide gene cassette such as DTA to achieve global ablation of microglia regardless of their states. To test if our Clec7a-CreER^T2^ line can be used to perform state-specific depletion, we crossed the driver line to Cre-dependent DTA and bred both loci to homozygosity to increase efficiency. Animals were placed on either a cuprizone only diet (control condition), or a combined cuprizone and tamoxifen diet (DAM-depletion condition) for five weeks (**Figure 8A**). Concurrent tamoxifen injections were administered in the depletion cohort to ensure that no new DAM were formed from the homeostatic pool or the residual DAM. We found almost complete absence of CLEC7A^+^ microglia during demyelination in the DAM-depletion group compared to the control group (**Figures 8B, 8C**). To continue ablation during remyelination, control animals were switched to a regular diet, and DAM-depletion animals were switched to a tamoxifen only diet with continued tamoxifen injections (**Figure 8A**). We observed a sustained suppression of CLEC7A^+^ microglia in our tamoxifen treated animals during remyelination (**Figure 8B**). In the absence of DTA, tamoxifen diets or injections had no overt negative effects on the formation of CLEC7A^+^ microglia (DAM) (**Figure 4H and Figure S6**), and therefore the lack of DAM in the depletion group was due to cell ablation but not tamoxifen treatment itself. Importantly, IBA1^+^CLEC7A^−^ microglia were still present in both gray and white matter regions, demonstrating that our depletion strategy was specific to DAM, while preserving homeostatic microglia.

**Figure 8.**
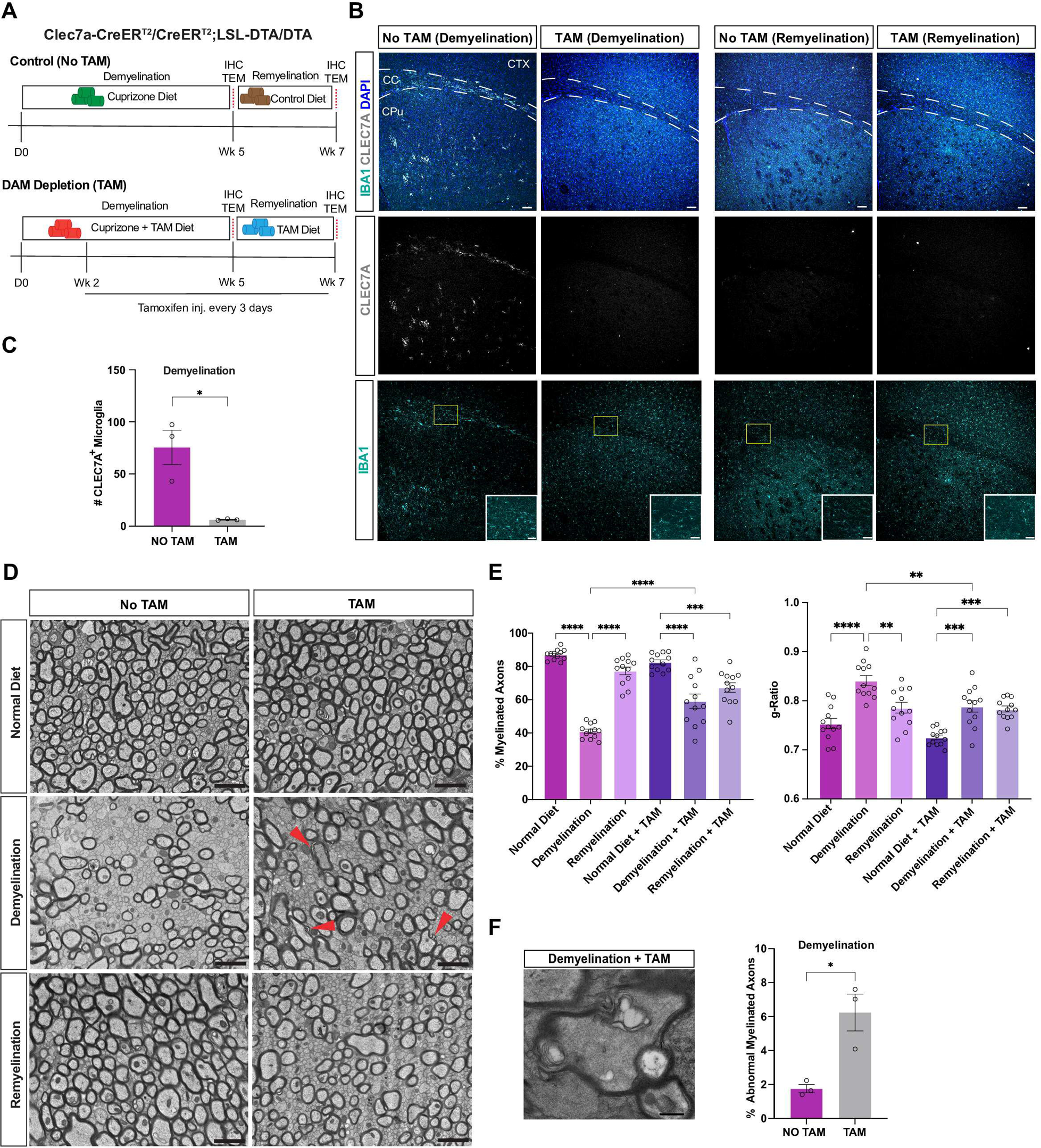
DAM depletion shows their protective roles for the recovery of cuprizone-induced demyelination. **(A)** Schematic of experimental design for control (top) and DAM depletion (bottom) models during demyelination and remyelination. **(B)** Representative immunostaining images showing CLEC7A^+^ microglia during demyelination (left two panels) and remyelination (right two panels) in control (No TAM) and DAM depletion (TAM) models. Scale bar = 100um. Boxed insert showing ramified IBA1^+^ microglia in CC. Scale bar = 50um. CC, corpus callosum; CTX, cortex; CPu, caudate putamen. **(C)** Quantification of CLEC7A^+^ microglia in the corpus callosum during demyelination in control and DAM depletion models. n = 3. Student’s T-test, * *p* < 0.05. Error bar represents +/− SEM. **(D)** Transmission electron microscopy images of axons in the corpus callosum during demyelination and remyelination in control (No TAM) and DAM depletion (TAM) models. Arrowheads point to axons with abnormal myelin patterns. Scale bar = 2um. **(E)** Quantification for the percentage of myelinated axons and g-ratio in the corpus callosum from Clec7a-CreER^T2^;DTA mice fed with normal diet, or mice at demyelination or remyelination stages, with or without tamoxifen. n = 12. One-way ANOVA with Tukey’s multiple comparisons test. ns: not significant, *** *p* < 0.001, **** *p* < 0.0001. Error bar represents +/− SEM. **(F)** Representative image (left) and quantification (right) of axons with abnormal myelin patterns during demyelination. Scale bar = 500nm. n = 3. Student’s T-test, * *p* < 0.05. Error bar represents +/− SEM. **See also Figure S6.**

Being able to specifically deplete DAM provided an opportunity to pinpoint their roles during demyelination and remyelination. We performed transmission electron microscopy (TEM) to examine the effect of DAM depletion on myelination in the cuprizone model. We observed that a higher number of axons still had myelin attached to them during demyelination upon DAM depletion compared to the control group, and these axons also displayed lower average g-ratio, indicative of thicker myelin, in the treatment group (**Figures 8D, 8E**). DAM removal seemed to cause less severe demyelination by these measurements; however, there was a higher frequency of axons wrapped by abnormal myelin (**Figures 8D, 8F**). Interestingly, we found that remyelination was less complete in the depletion group compared to control animals (**Figures 8D, 8E**). Together, these results support the model that the DAM state plays an active role in damaged myelin removal, which is required for myelin repair following cuprizone-induced demyelination.

## DISCUSSION

Microglia are highly dynamic cells with a range of functional roles in development and disease. Previous studies suggest that these diverse functions may be regulated by distinct microglial states, defined by the expression of unique gene casettes.^36,61^ However, the inability to label and manipulate microglia in a state-specific manner has made it challenging to make definitive claims about the biological implications of microglial heterogeneity. In this study, we created a new inducible genetic tool, Clec7a-CreER^T2^, and characterized its labeling efficiency and specificity for a series of microglial states, including PAM during early postnatal development, and DAM across several disease models. We demonstrated three potential applications: (1) acute isolation of these microglial subpopulations for transcriptomic analysis; (2) fate mapping to track dynamic changes of microglial states; (3) state-specific depletion for functional studies. These experiments allowed us to define converging and context-dependent gene pathways in similar microglial states, and uncovered the plasticity and protective roles of disease-associated microglia in a model of white matter disease.

Numerous microglial states have been identified through single-cell genomics, which has outpaced our understanding of their functional relevance.^32,34–36^ The generation of new driver lines such as Clec7a-CreER^T2^ holds the promise to bridge the major gap between molecular characterizations and biological function. Our CreER reporter faithfully labels up to 90% PAM during development, and DAM-like cells with 60-85% efficiency across the tested disease models. The range of labeling efficiency in these contexts, which is on par with the existing microglial driver lines,^46,47^ may be attributed to differences in *Clec7a* expression and tamoxifen dosing. We have not observed spontaneous recombination without tamoxifen, and homozygosity of the driver does not appear to affect *Clec7a* gene function or cause haploinsufficiency. Therefore, if necessary, homozygous Clec7a-creER^T2^ may be used in combination with optimized tamoxifen regimens to maximize labeling efficiency.

Differences in microglial isolation protocols and sequencing platforms often affect microglial transcriptomics and interpretation of sequencing results.^36,56^ Bioinformatic tools can remove batch effects to certain extent,^62^ but cross-datasets integration is still challenging, which partially led to a poorly defined nomenclature of microglial states.^36^ By targeted enrichment of specific microglial subpopulations, we benchmarked a systematic comparison of PAM and various DAM-like substates using a standardized experimental pipeline.

Through this analysis, we described 11 upregulated genes shared among these microglial states as a core signature, and meanwhile demonstrated context-dependent gene activation. For example, *Spp1*, *Cst7* and *Lgals3*, which have been grossly considered as part of the PAM/DAM gene cassette, seem to have biases in their levels of upregulation among PAM, AD-DAM and EAE-DAM, respectively. GO term analysis further revealed extensive differences between PAM and DAM, where PAM are more associated with biosynthetic and metabolic pathways and DAM tend to enrich immune response terms. While this may not be surprising as these microglial states appear in opposing contexts, it raises questions about the functional significance of the shared signature.

A lack of such knowledge also contributes to the controversy on the use of DAM as an umbrella term for microglia responding in pathological conditions.^36^ We use this nomenclature here solely to simplify the narrative. Contrasting DAM from 5xFAD and EAE showed that AD-DAM upregulated pathways related to complement signaling, astrocyte activation and T cell proliferation, which have all been implicated in AD pathology,^6,63,64^ whereas EAE-DAM were involved in the recruitment and regulation of various peripheral immune cells. Future work should address which and how environmental signals drive both the unique and shared gene signatures.

A key question in the field has been the extent to which reactive microglial states are plastic and reversible. By genetic fate mapping in the cuprizone model, we provided an example for the transcriptomic and morphological plasticity of DAM-like cells in the context of white matter injury. The same microglial cell can be activated and subsequently return to homeostasis, which is presumably competent for additional rounds of activation. Given that microglia are long lived,^65–68^ functional consequences of such state alternation deserve further investigation. Future studies should also examine the plasticity of microglial states in diseases that are progressive in nature.

Traditional methods of microglia depletion lack state-specificity. In addition, certain microglial subpopulations are resistant to the commonly used PLX drugs, possibly due to the downregulation of *Csf1r*.^58,69,70^ As a proof of principle, we used Clec7a-CreER^T2^;LSL-DTA to demonstrate near complete depletion of DAM in the cuprizone model. This allowed us to pinpoint their function, which is responsible for removing damaged myelin to facilitate efficient remyelination. These results are consistent with previous studies using *Trem2* knockout mice in the same model.^71^ While we focused on cell ablation, the Clec7a-CreER^T2^ mouse can also be used to perform state-specific gene knockouts.

Lastly, given that *Clec7a* is often upregulated in many other microglial states,^72^ we believe that this highly versatile tool will be useful for parsing out the distinct roles of microglia in development, aging, and a variety of neurodegenerative diseases, such as amyotrophic lateral sclerosis, frontotemporal dementia, Parkinson’s disease and beyond.

### Limitations of the study

Although the Clec7a-CreER^T2^ driver specifically targets microglial subpopulations in the CNS parenchyma under normal conditions, it also labels, albeit at lower levels, CNS border-associated macrophages (BAM) and certain peripheral myeloid cells. This should be taken into account particularly for models that involve peripheral immune infiltration. Due to this limitation, we were unable to distinguish between resident DAM and infiltrating myeloid cells that might have acquired the DAM signature in EAE.^73^ For experiments that require higher levels of specificity, generating wildtype bone marrow chimeras in the Clec7a-CreER^T2^ mouse may be helpful. Another caveat is that *Clec7a^+^* microglia may include more than one state. This is exemplified in EAE, where labeled microglia fell into the MHCII^+^ cluster and classical DAM clusters. Staining with multiple markers or ideally single cell profiling can help determine the composition of targeted cells and mitigate the risk of data misinterpretation. In the future, development of split Cre models will allow further dissection of co-labeled microglial substates and BAM.

## ACKNOWLEDGEMENTS

We thank the Genome Technology Access Center (GTAC) at the McDonnell Genome Institute at Washington University School of Medicine for help with sequencing. The Center is partially supported by NCI Cancer Center Support Grant #P30 CA91842 to the Siteman Cancer Center from the National Center for Research Resources (NCRR), a component of the National Institutes of Health (NIH), and NIH Roadmap for Medical Research. We also thank Jennifer Ponce at GTAC for providing library preparation liquid handlers and Xiaoxia Cui at Genome Engineering & Stem Cell Center for providing the FACS machine. We thank Lu Sun, Ye Zhang, Hui Zong, Gregory Wu, Anne Cross, and the ZYGL group for generous discussion on the project. We thank Darsh Singhania for his technical support. N.S. is supported by the Bursky Center for Human Immunology and Immunotherapy Programs. G.Y. is supported by the NIH (R01MH110504 and U19NS123719). N.A. is supported by American Heart Association (#23PRE1025832). Q.L. is supported by the Whitehall Foundation (2021-08-003), BIG Center at WashU, ICTS, the Hope Center for Neurological Disorders, the McDonnell Center for Cellular and Molecular Neurobiology and NIH (R01AG078512).

## CONTACT FOR REAGENT AND RESOURCE SHARING

Further information and requests for resources and reagents should be directed to and will be fulfilled by the Lead Contact, Qingyun Li

## AUTHOR CONTRIBUTIONS

Conceptualization, K.B., N.A., and Q.L.; Methodology, K.B., N.A., Z.C., M.K., L.Z., J.Y., J.R., J.P., L.S., W.B.; Software, K.B, Z.C., Q.Y.; Investigation, K.B., N.A., Z.C.; Formal Analysis, K.B., N.A., Z.C.; Writing – Original Draft, K.B., N.A., Z.C.; Writing – Review & Editing, K.B., Q.L.; Funding Acquisition, Q.L.; Resources, M.K., J.K., G.Y., J.H., N.S., M.C.; Supervision, Q.L., G. Y., J.K., M.C.

## DECLARATION OF INTERESTS

The authors declare no competing interest

## METHODS

### Mice

All experiments were performed under the approval of the Institutional Animal Care and Use Committee at Washington University in St. Louis. Mice were housed under pathogen-free conditions, with controlled temperature and humidity, a 12-hour light/dark cycle, and no more than five mice per cage. Rodent chow and water were provided *ad libitum*. Cages and bedding were changed weekly.

### Generation of Clec7a-P2A-iCreER^T2^ mice by CRISPR-Cas9

Clec7a-P2A-iCreER^T2^ mice were generated by CRISPR-Cas9 knock in. Specifically, the gRNA was designed by the CRISPR design tool (http://crispr.mit.edu) to target the region between exon 5 and 3’ UTR of the *Clec7a* gene locus. On-target activity of gRNA was screened using UCA^TM^ (Universal CRISPR Activity Assay), a sgRNA activity detection system developed by Biocytogen. To generate T7-Cas9/sgRNA products, T7 RNA polymerase was added to the Cas9 or sgRNA templates *in vitro* by PCR amplification. After purification, products were used as the template for *in vitro* transcription using the MEGAshortscript T7 kit (Life Technologies) according to the kit protocol. To test that the P2A-iCreER^T2^ was precisely inserted before stop codon of the *Clec7a* gene, a circular donor vector was used. The gene targeting vector containing P2A-iCreER^T2^ and 2 homology arms of left (1009 bp) and right (2094 bp) each was used as a template to repair the DSBs generated by Cas9/sgRNA. To test *in vivo*, Cas9 mRNA and sgRNA were mixed and co-injected into the cytoplasm of one-cell stage fertilized eggs collected from C57BL/6N females, which were transferred to psuedopregnant females. DNA samples from F1 progeny were tested for targeted insertion of P2A-iCreER^T2^ in *Clec7a* gene by southern blot analysis. This line was crossed to the C57BL/6J background.

### Mouse lines

C57BL/6J were used as wildtype mice. Cx3cr1^GFP^ (stock #005582-JAX), Rosa-LSL-tdTomato (Ai14) (stock #007914-JAX), Rosa-LSL-DTA (stock #009669-JAX) mice were purchased from Jackson Laboratories. 5xFAD mice were obtained from the MMRRC (stock #034848-JAX). CD11c-CreERT mice were purchased from the European Mouse Mutant Archive (EMMA: 09004). Female mice tend to display a more severe clinical response in EAE model and were chosen for sequencing analysis. To control for any sex specific transcriptomic differences, and standardize cross-condition comparison, female mice were also used for all other sequencing experiments. For non-sequencing related studies, a combination of male and female mice were used.

### Experimental Models and Tamoxifen delivery

For P7 pups, a single 10uL injection of a 10mg/mL tamoxifen solution prepared in oil was subcutaneously injected into each mouse at either P4 or P5. For 5xFAD labeling, Clec7a-CreER^Τ2^;LSL-tdTomato;5xFAD mice were raised to 3 months of age and 200uL of a 20mg/mL tamoxifen solution was injected intraperitoneally. For cuprizone labeling, Clec7a-CreER^T2^;LSL-tdTomato mice were aged to 2-3 months and were administered 0.2% cuprizone for 5 weeks, and switched to a control diet for 2 weeks of remyelination. Cuprizone treated animals were administered 200uL of 20mg/mL tamoxifen solution intraperitoneally. All adult mice for 5xFAD and cuprizone labeling experiments received injections every other day for 3 days, followed by 3 daily injections and sacrifice. For EAE experiment, Clec7a-CreER^T2^;LSL-tdTomato mice were aged to 2-4 months and were immunized with 200ug of MOG (35-55), 1mg/mL of complete Freund’s adjuvant supplemented with BD Difco Mycobacterium Tuberculosis H37 Ra Dessicated and 300ng pertussis toxin. Mice used for EAE sequencing experiment received 200ng pertussis toxin. Immunized mice received a single 200uL injection of 20mg/mL tamoxifen solution at peak disease score and were sacrificed 48hr later. For DAM depletion experiment, Clec7a-CreER^T2^;LSL-DTA mice were aged to 2-4 months and were fed diets containing 0.2% cuprizone, 0.2% cuprizone + 400ppm tamoxifen, 400ppm tamoxifen, or control diets for 5 weeks, and switched to control diets or 400ppm tamoxifen diets for 2 weeks for remyelination experiments. Mice used for the DAM depletion group received 200uL injections of 20mg/mL tamoxifen solution every 3 days for duration of the experiment.

### Experimental Autoimmune Encephalomyelitis Clinical Scoring

One week after immunization, animals were scored daily for locomotor deficits. Score assessments were given as follows: 0 – no behavior deficits; 0.5 – straining of tail base; 1 – limp tail; 1.5 – abnormal gait; 2 – tail base lowered while walking; 2.5 – dependence on one hind limb; 3 – paralysis of both hind limbs; 4 – hind and fore limb paralysis; 5 – death.

### Immunohistochemistry

Mice were euthanized with a ketamine and xylazine cocktail prepared in saline. Mice were then transcardially perfused with PBS followed by 4% PFA. Brains were extracted and fixed in 4% PFA overnight at 4°C, then cryopreserved in 30% sucrose for 48 hours at 4°C. Brains were embedded in optimal cutting temperature medium (Tissue-Tek) and manually sectioned on a cryostat (Leica). For dura, leptomeninges, and choroid plexus dissection, separate mice were transcardially perfused with PBS. Duras were peeled from the dorsal skull caps, leptomeninges were peeled from the surface of dissected brains, and choroid plexuses were dissected from the lateral ventricles. Dissected tissues were fixed for 1 hour in 4% PFA, then washed with PBS and mounted onto glass slides for staining.

Free-floating sections and flat-mounted tissues were blocked and permeabilized in PBS with 10% serum and 0.2% Triton X-100 for 15 minutes at room temperature. Tissues were incubated in primary antibodies diluted in PBS with 1% donkey serum and 0.2% Triton X-100 overnight at 4°C. Tissues were washed three times, incubated in secondary antibodies diluted in PBS with 1% serum and 0.2% Triton X-100 for 2 hours at room temperature, then washed another three times. For brain sections stained with Amylo-Glo (Biosensis TR-300), tissue was next incubated in a 1:500 solution of Amylo-Glo diluted in PBS for 20 minutes then washed three times. For sections stained with FluoroMyelin, tissues sections were incubated in 1:100 of FluoroMyelin diluted in PBS for 30 minutes then washed three times. Free-floating brain sections were transferred onto glass slides. Slides were coverslipped using Vectashield without DAPI for Amylo-Glo tissues (Vector Laboratories H-1000) or Vectashield with DAPI for all other samples (Vector Laboratories H-1200). Images were acquired using a Nikon A1R confocal microscope.

The following primary antibodies were used: Goat polyclonal anti-IBA1 (Abcam ab5076, 1:500); Rat neutralizing monoclonal anti-mDECTIN-1-IgG (clone R1-4E4) (Invivogen mabg-mdect, 1:30); Rat monoclonal anti-CD206 (clone MR5D3) (BioRad, MCA2235); Armenian hamster monoclonal anti-CD31 (clone 2H8) (Invitrogen, MA3105); Rabbit anti-Olig2 (EMD Milipore Sigma AB9610, 1:1000); Hamster anti-CD11c (Bio-Rad MCA1369, 1:40); Rabbit anti-CD68 (Abcam 283654, 1:100); Chicken anti-Myelin Basic Protein (AVES 0200 1:500); Mouse anti-β-Amyloid, 1-16 (clone 6E10) (Biolegend, 803013, 1:200).

The following secondary antibodies were used: Donkey anti-goat IgG (H+L) cross-adsorbed, Alexa Fluor 488 (Thermo Fisher Scientific, 11055, 1:500); Donkey anti-goat IgG (H+L) AffiniPure, Alexa Fluor 647 (Jackson Immuno Research Laboratories 705-605-147, 1:500); Donkey anti-rat IgG (H+L) AffiniPure, Alexa Fluor 647 (Jackson Immuno Research Laboratories 712-605-153, 1:200); Alexa Fluor 488 AffiniPure Goat Anti-Armenian Hamster IgG (H+L) (Jackson Immuno Research Laboratories, 127-545-160); Alexa Fluor 647 AffiniPure Goat Anti-Rat IgG (H+L) (Jackson Immuno Research Laboratories, 112-605-003); Alexa Fluor 488 Donkey anti-rabbit (Thermo Fisher Scientific A-21206, 1:1000); Alexa Fluor 647 Donkey anti-Rabbit (Thermo Fisher Scientific, A-31573, 1:200); Alexa Fluor 647 Goat anti-Rabbit IgG (H+L) (Thermo Fisher Scientific A-21245, 1:200).

### EdU (5-ethynyl-2’-deoxyuridine) labeling

EdU powder was dissolved in 1XPBS by agitating and heating mixture to 37-40°C, 30 minutes before injection. EdU solution was administered intraperitoneally at a dose of 50mg/kg in 100uL every three days during two weeks of remyelination. Standard immunohistochemistry procedure was performed prior to EdU development. To develop EdU stain, sections were first permeabilized in 0.5% PBST for 30 mins. Sections were then incubated in an EdU development cocktail for 30 mins at room temperature and washed three times in 0.5% PBST. EdU development cocktail was prepared according to the following concentrations and order listed, no more than 15 minutes before incubation: 75.75% 1XTBS (pH 7.6), 4% 100mM copper sulfate in water, 0.25% 2mM sulfo-Cy5 azide in DMSO, and 20% 500mM sodium ascorbate in water.

### Confocal Image analysis

To quantify labeling efficiency and specificity in Cre reporter lines, manual counting of CLEC7A^+^ and tdTomato^+^ cells and quantification of double-positive cells were performed in Fiji, using the Cell Counter plugin. Microglia morphology was quantified via manual counting of primary branches and terminal branch points in Fiji. At least 15 cells per condition were quantified. 3D reconstructions of microglia were generated with the Imaris software. To quantify the distance of tdTomato^+^ microglia to plaques, the total number of tdTomato^+^ microglia within a field of view was counted, and the distance of each cell body to the nearest plaque was manually measured. Data represent averages for three mice. At least 3 images per mouse, each containing an average of 73 cells, were quantified. The percentage of the tdTomato^+^ area divided by the total area in the spinal cord was quantified using Fiji. At least 3 mice and 3 sections per mouse were quantified for each score bin.

### Phagocytosis Assay

Acute brain slices (250um thickness) from Clec7a-CreER^T2^ heterozygous, homozygous and WT P7 littermates were cut using a vibratome. Immediately after cutting, slices were incubated in slice culture media (65%MEM, 10% FBS, 25% HBSS, 6.5 mg/mL Glucose, 2mM Glutamine, 1% Penicilin/Streptomycin) for 1 hour in an incubator (37°C, 5% carbon dioxide). After incubation, sections were further incubated in zymosan coated pHrodo beads (1mg/mL, Invitrogen, P35364), making sure to cover corpus callosum region, for 4 hours in the incubator. Finally, sections were fixed in 4% PFA for 2 hours. Standard immunohistochemistry protocol and quantification was performed as previously described.

### Transmission Electron Microscopy and Myelin Quantification

Animals were transcardially perfused with 20mL of PBS. The left hemisphere was dropped fixed in 4% PFA for immunohistochemistry analysis. A 1mm slice cut from the medial side of the right hemisphere was fixed in 2% paraformaldehyde/2.5% glutaraldehyde (Ted Pella Inc., Redding, CA) in 100mM cacodylate buffer, pH 7.2 for 2 hours at room temperature and then overnight at 4°C. Samples were washed in cacodylate buffer and postfixed in 1% osmium tetroxide (Ted Pella Inc.) for 1 hour. Samples were then rinsed extensively in dH_2_0 prior to en bloc staining with 1% aqueous uranyl acetate (Ted Pella Inc.) for 1 hour. Following several rinses in dH_2_0, samples were dehydrated in a graded series of ethanol and embedded in Eponate 12 resin (Ted Pella Inc.). Sections of 95nm were cut with a Leica Ultracut UCT ultramicrotome (Leica Microsystems Inc., Bannockburn, IL), stained with uranyl acetate and lead citrate, and viewed on a JEOL 1200 EX II transmission electron microscope (JEOL USA Inc., Peabody, MA) equipped with an AMT 8 megapixel digital camera (Advanced Microscopy Techniques, Woburn, MA). 5000X and 15000X images of corpus callosum were taken for analysis. G-ratio (diameter of axon divided by the diameter of axon plus myelin) was quantified using MyelTracer Software [https://github.com/HarrisonAllen/MyelTracer].

### Cell isolations for flow cytometry

Mice were injected with 10% Euthasol and transcardially perfused with PBS. Brains and livers were harvested from each mouse. Duras were peeled from the reserved dorsal skull caps, leptomeninges were peeled from the surface of the dissected brains, and choroid plexuses were dissected from the lateral ventricles. Tissues were then transferred to digestion buffer (DMEM with 1 mg/mL Collagenase VIII (Sigma Aldrich), 0.5 mg/mL DNase I (Sigma-Aldrich), and 2% FBS) and mechanically dissociated. Tissues were incubated at 37°C for either 15 minutes (meninges and choroid plexus) or 30 minutes (liver and brain). Samples were triturated with a P1000 pipette 5 times, passed through a 70um cell strainer, and centrifuged at 500*g* for 5 minutes at 4°C. Cells were resuspended in an equal volume of DMEM containing 10% FBS to inactivate enzymatic digestion. After pelleting at 500*g* for 5 min, samples from the meninges, choroid plexus, and liver were resuspend in FACS buffer (PBS with 2% BSA and 1mM EDTA) and kept at 4°C for staining. To remove myelin, brain samples were resuspended in 9mL of 22% BSA in a 1:1 solution of PBS to DMEM. Brain samples were then centrifuged at 1000*g* for 10 min at 4°C, with a brake of 3, and the upper myelin-containing layer was aspirated along with the remaining BSA. Samples were resuspended in FACS buffer and kept at 4°C for staining. For blood samples, blood was collected prior to perfusion and centrifuged at 500*g* for 5 minutes at 4°C. Red blood cells were lysed by resuspension in 1mL of ACK lysis buffer (Quality Biological) for 10 minutes at room temperature, followed by addition of 2mL of PBS to inactivate the ACK lysis buffer. Samples were pelleted, resuspended in FACS buffer, and kept at 4°C for staining.

For staining of meninges, choroid plexus and peripheral organs, cells were transferred to a 96-well V-bottom plate and pelleted (500*g* for 5 minutes at 4°C). Viability staining was performed using Zombie NIR for 10 minutes at room temperature (1:500 in PBS, BioLegend). Cells were pelleted and resuspended in anti-CD16/32 antibody (1:200, BioLegend) diluted in FACS buffer to block Fc receptor binding. Cells were incubated in fluorophore-conjugated antibodies diluted in FACS buffer for 10 minutes at room temperature. Flow cytometry was performed using an Aurora spectral flow cytometer (Cytek Biosciences). Data were analyzed with FlowJo (version 10, BD Biosciences).

The following antibodies were used: BV750 Rat Anti-Mouse CD45 (BD, 746947); BUV661 Rat Anti-Mouse Ly-6G (BD, 741587); BUV563 Rat Anti-CD11b (BD, 741242); Brilliant Violet 711 anti-mouse CD64 (FcγRI) Antibody (BioLegend, 139311); Alexa Fluor 700 anti-mouse F4/80 Antibody (BioLegend, 123130); Alexa Fluor 488 anti-mouse CD206 (MMR) Antibody (BioLegend, 141710); Brilliant Violet 510 anti-mouse I-A/I-E Antibody (BioLegend, 107635); BUV737 Hamster Anti-Mouse CD11c (BD, 749039); BV480 Rat Anti-Mouse CD19 (BD, 566107); PerCP/Cyanine5.5 anti-mouse Ly-6C Antibody (BioLegend, 128012); BUV805 Hamster Anti-Mouse TCR β Chain (BD, 748405); PE/Cyanine7 anti-mouse NK-1.1 Antibody (BioLegend, 108714); BD Horizon BUV395 Rat Anti-Mouse CD4 (BD, 563790); CD8a Monoclonal Antibody (53-6.7), Alexa Fluor 532, eBioscience (Invitrogen, 58-0081-80); BV711 Rat Anti-Mouse CD45 (Biolegend, 103147); BV421 Rat Anti-Mouse CD45 (Biolegend, 101236).

### Microglia isolation for FACS from brain and spinal cord tissue

Mice were injected with ketamine and transcardially perfused with PBS. Microglia cells were isolated using a previously published protocol.^57^ Briefly, animals were transcardially perfused with 20 mL of PBS. Brains or spinal cords were finely chopped using a blade and were further homogenized using a douncer and piston. Homogenized tissue solution was filtered using a 70um cell strainer, centrifuged (500*g* for 5 minutes at 4°C) and resuspended in MACS buffer (0.5% BSA, 2mM EDTA, in 1x PBS). Tissue solution was then incubated in myelin removal beads (Miltenyi Biotec,130-096-733) and filtered through LD columns (Miltenyi Biotec, 130-042-901). The single cell suspension was centrifuged and resuspended in PBS for LIVE/DEAD staining for 10 minutes (1:1000, Life Technologies, L34970). After LIVE/DEAD stain was washed, cells were resuspended in FACS buffer and incubated in FC Block for 5 minutes (1:60, BD Pharmingen, 553142) and primary antibodies for 10 minutes. Antibodies were washed, cells were resuspended in FACS buffer and RNase inhibitor, and kept at 4^°^C for sorting. Flow cytometry was performed using the SONY sorter (SH800). Microglia were classified CD45^intermediate^CD11b^+^ cells, gated on live singlets. Single tdTomato^+^ or tdTomato^−^ cells from each animal were sorted into 96-well plates previously prepared with lysis buffer. Once plates were fully sorted, they were immediately vortexed, centrifuged and stored in −80°C until downstream library preparation and sequencing analysis was performed.

### Lysis Buffer Preparation for single cell RNA sequencing plates

Each lysis plate well contained 4uL of lysis buffer (4U Recombinant RNase Inhibitor (Takara Bio 2313B), 0.05% Triton X-100, 2.5mM dNTP mix (Thermo Fisher Scientific R0192), 2.5uM Oligo-dT30VN (5′-AAGCAGTGGTATCAACGCAGAGTACT30VN-3′. Plates were stored at −80°C freezer until use.

### scRNA-seq library preparation

Sorted microglia plates were prepared for scRNA-sequencing following the previously published Smart-seq2 protocol.^57^ Briefly, plates were thawed and primer annealing was performed at 72°C for 3 minutes. Reverse transcription was performed by adding 6uL of reverse transcription mixture (95U SMARTScribe Reverse Transcriptase (100U/uL, Clontech 639538), 10U RNase inhibitor (40U/uL), 1XFirst-Strand buffer, 5mM DTT, 1M Betaine, 6mM MgCl2, 1uM TSO (Rnase free HPLC purified) to each well (PCR protocol - at 42°C for 90 min, followed by 70°C, 5 min). DNA preamplification was done by adding 15 uL of PCR amplification mix (1X KAPA HIFI Hotstart Master Mix (Kapa Biosciences KK2602), 0.1uM ISPCR Oligo (AAGCAGTGGTATCAACGCAGAGT), 0.56U Lambda Exonuclease (5U/uL, New England BioLabs M0262S) to each well (PCR protocol - (1) 37°C 30 min; (2)95°C 3 min; (3) 23 cycles of 98°C 20 s, 67°C 15 s, 72°C 4 min; (4) 72°C 5 min). cDNA was cleaned using PCRClean DX beads (0.7:1 ratio, Aline C-1003-50) and resuspended in 20uL EB buffer.

Nextera XT DNA Sample Prep Kit (Illumina FC-131-1096) was used at 1/10 of recommendation volume, with the help of a Mosquito HTS robot for liquid transfer. Specifically, tagmentation was done in 1.6uL (1.2uL Tagment enzyme mix, 0.4uL cDNA) at 55°C, 10 min. To stop the reaction, neutralization buffer was added 0.4uL per well and incubated at room temperature for 5 min. Then 0.8uL Illumina 10-bp dual indexes (0.4uL each, 5uM) and 1.2uL PCR master mix were added to amplify whole transcriptomes using the following program: (1) 72°C 3 min; (2) 95°C 30 s; (3) 10 cycles of 95°C 10 s, 55°C 30 s, 72°C 1 min; (4) 72°C 5 min. Libraries from a single 384 plate were pooled together in an Eppendorf tube and purified twice with PCRClean DX beads. The quality and concentrations of the final mixed libraries were measured with Bioanalyzer and Qubit, respectively, before Illumina Nova sequencing. To maximize PCR amplification efficiency for cuprizone demyelination samples, biotinylated-oligo-dT30VN oligo and biotinylated-TSO oligo were used in place of Oligo-dT30VN and TSO respectively, as described in an enhanced Smart-seq2 protocol.^74^

### Processing of scRNA-seq raw data

The alignment of scRNA-seq raw data was conducted using the same pipeline as previously described.^28^ Specifically, Prinseq^75^ was utilized to filter sequencing reads shorter than 30 bp (-min_len 30), trim the first 10 bp at the 5′ end (-trim_left 10) of the reads, trim reads with low quality from the 3′ end (-trim_qual_right 25), and remove low complexity reads (-lc_method entropy, -lc_threshold 65). Afterward, Trim Galore was deployed to remove the Nextera adapters (–stringency 1), followed by STAR^76^ to align the remaining reads to the mm10 genome using the settings: – outFilterType BySJout,–outFilterMultimapNmax 20,–alignSJoverhangMin 8,– alignSJDBoverhangMin 1,–outFilterMismatchNmax 999,–outFilterMismatchNoverLmax 0.04,–alignIntronMin 20,–alignIntronMax 1000000,–alignMatesGapMax 1000000,– outSAMstrandField intronMotif. Then Picard was employed to remove the duplicate reads (VALIDATION_STRINGENCY = LENIENT, REMOVE_DUPLICATES = true). Ultimately, the aligned reads were converted into counts for each gene by using HTSeq (-m intersection-nonempty, -s no).^77^

### Clustering analysis of scRNA-seq data

The Seurat package was utilized to perform unsupervised clustering analysis on scRNA-seq data.^78^ Briefly, gene counts for cells that passed QC thresholds (percent.ribo < 0.08, total_counts >1e+5, and number_of_expressed_genes > 300) were normalized to the total expression and log-transformed, and highly variable genes were detected. With the top 2000 highly variable genes as input, principal component analysis (PCA) was conducted on the scaled data, and the top components were utilized to compute the distance metric. This distance metric then underwent unsupervised cell-clustering analysis. Based on the initial clustering results, we further subsisted the microglia-related clusters for finer clustering analysis. The results were visualized in a low-dimension projection using Uniform Manifold Approximation and Projection (UMAP).^79^

### DE gene analysis

To identify the differentially expressed genes (DEGs) of each cluster, we conducted a DEG analysis (one versus rest) on the scRNA-seq data, which is done by using the FindAllMarkers function in the Seurat package. In our case, this function executed Wilcoxon rank sum tests with following settings: logfc.threshold = 0.25, min.pct = 0.1. Additionally, DEG analysis was also performed separately for each condition (P7, 5xFAD, or EAE). In this context, within each condition (P7, 5xFAD, or EAE), the DEG analysis compared DAM (or PAM) cells against the corresponding homeostatic microglia, using the same statistical test method and settings as mentioned above. A threshold of significance was set at p-value_adj < 0.05 after applying the Benjamini-Hochberg (BH) procedure for multiple test correction.

### DEG comparison between conditions

To compare the gene expression changes of PAM/DAM cells in any two different conditions (P7 vs. 5xFAD, P7 vs. EAE, and 5xFAD vs. EAE), a z-score was assigned to each gene in each condition, signifying the significance of the gene’s expression change from homeostatic levels within that condition. Subsequently, a scatter plot was created to compare the z-scores of all genes between any two conditions. The z-score for each gene was obtained through differential gene expression (DEG) analysis, specifically using Wilcoxon rank-sum tests with the following settings: logfc.threshold = 0, min.pct = 0. This analysis compared PAM/DAM cells to the corresponding homeostatic microglia in each condition. Each *p*-value was then transformed into a standardized score, z-score.

### Similarity Score Calculation

TySim^80^ was utilized to quantify to what extent each individual cell is similar to a specific target cell type (e.g., PAM or DAM). A cell is considered similar to a target cell type at the transcriptional level if it expresses the signatures of the target cell type that would be expected by random chance with statistical significance. Given the signatures of target cell type and the scRNA-seq data to be tested, TySim conducts statistical test to access the similarity levels. Briefly, TySim separates the scRNA-seq data into binary part and non-zero part and handles them separately so as to mitigate the impact of drop-out effects in scRNA-seq datasets. In each part of data, TySim takes into account artifact factors that can jointly influence observed expression values in scRNA-seq data, such as variations in sequencing depth across cells and heterogeneous preferences for different genes during sequencing. TySim considers the background expression attributable to these artifacts for each gene within each cell. These background expression levels are precisely estimated by systematically modeling both cell and gene factors embedded within the inputted scRNA-seq data, a process achieved by employing the Conditional Multifactorial Contingency (CMC) model. In this study, the signatures of PAM and DAM are obtained from previously published papers.^28,30^ The code of TySim is obtained from https://github.com/yu-lab-vt/CMC/tree/CMC-TySim.

### GO term analysis

The PANTHER Classification System was utilized to conduct the Gene Ontology (GO) term analysis.^81^ The inputs are the genes significantly upregulated in PAM/DAM compared to the corresponding homeostatic microglia within each condition. A statistical overrepresentation test was performed using Fisher’s Exact test, with default settings applied. Both fold enrichment values and the false discovery rate (expressed as −log10 FDR) for the enriched GO Biological Process (BP) terms were reported. The adjusted p-value was calculated through multiple test correction, utilizing the BH procedure.

### Quantification and Statistical Analysis

To quantify fluorescence intensity of Clec7a expression in P7 homozygous, heterozygous and wildtype littermates, representative tissue sections were selected and fluorescence intensity from the relevant channel was measured with Fiji by subtracting background signals from area integrated intensity. To quantify the numbers of CLEC7A^+^ microglia in corpus callosum of each genotype, cells on 50um sagittal brain sections were counted with Fiji (n = 3 sections for each genotype), and results were summarized as mean ± SEM. To do quantification in the phagocytosis assay, the total number of cells colabeled with beads from 3 randomly chosen fields on each 50um sagittal section were analyzed (5 sections per condition). The number of myelinated axons and abnormally myelinated axons were manually quantified using Fiji, Cell Counter plugin. Three mice were analyzed per group, and 4 sections per animal were used for quantification. To calculate statistical significance (p<0.05) for percent myelinated axons and g-ratio measurements, one-way ANOVA followed by Tukey’s multiple comparison post hoc test was performed on GraphPad prism. Summarized data were represented by mean +/− SEM. To calculate mean labeling efficiency, specificity, and morphology, 3 IHC sections (35um) from 3 animals were analyzed using Fiji and GraphPad Prism. Statistical significance (p < 0.05) for each measurement was calculated by either one-way ANOVA followed by Tukey’s multiple comparison post hoc test or Student’s T-test. Summarized data were represented by mean +/− SEM. To quantify percent of tdTomato area expressed in spinal cord of EAE, 3 sections from 3 mice were measured. Pearson correlation was calculated on GraphPad prism.

### Data and Software Availability

The accession numbers for the raw sequencing data reported in this paper will soon be available in a public repository.

## Supplemental Information

**Table S1.** DEGs comparing each microglia cluster to all the other clusters. **Related to Figure 5**.

**Table S2.** Genes upregulated in reactive microglial clusters (PAM or DAM) compared to condition-specific homeostatic clusters isolated from P7, 5xFAD and EAE conditions. **Related to Figure 6**.

**Table S3.** Pairwise comparisons for DEGs between P7, 5xFAD and EAE conditions. **Related to Figure 6**.

**Table S4.** Shared and unique GO terms associated DEGs from P7, 5xFAD and EAE conditions. **Related to Figure 6**

## Notes

### Competing Interest Statement

The authors have declared no competing interest.

