## Supplemental Figures for "An inducible genetic tool for tracking and manipulating specific microglial states in development and disease"

**Figure S1**

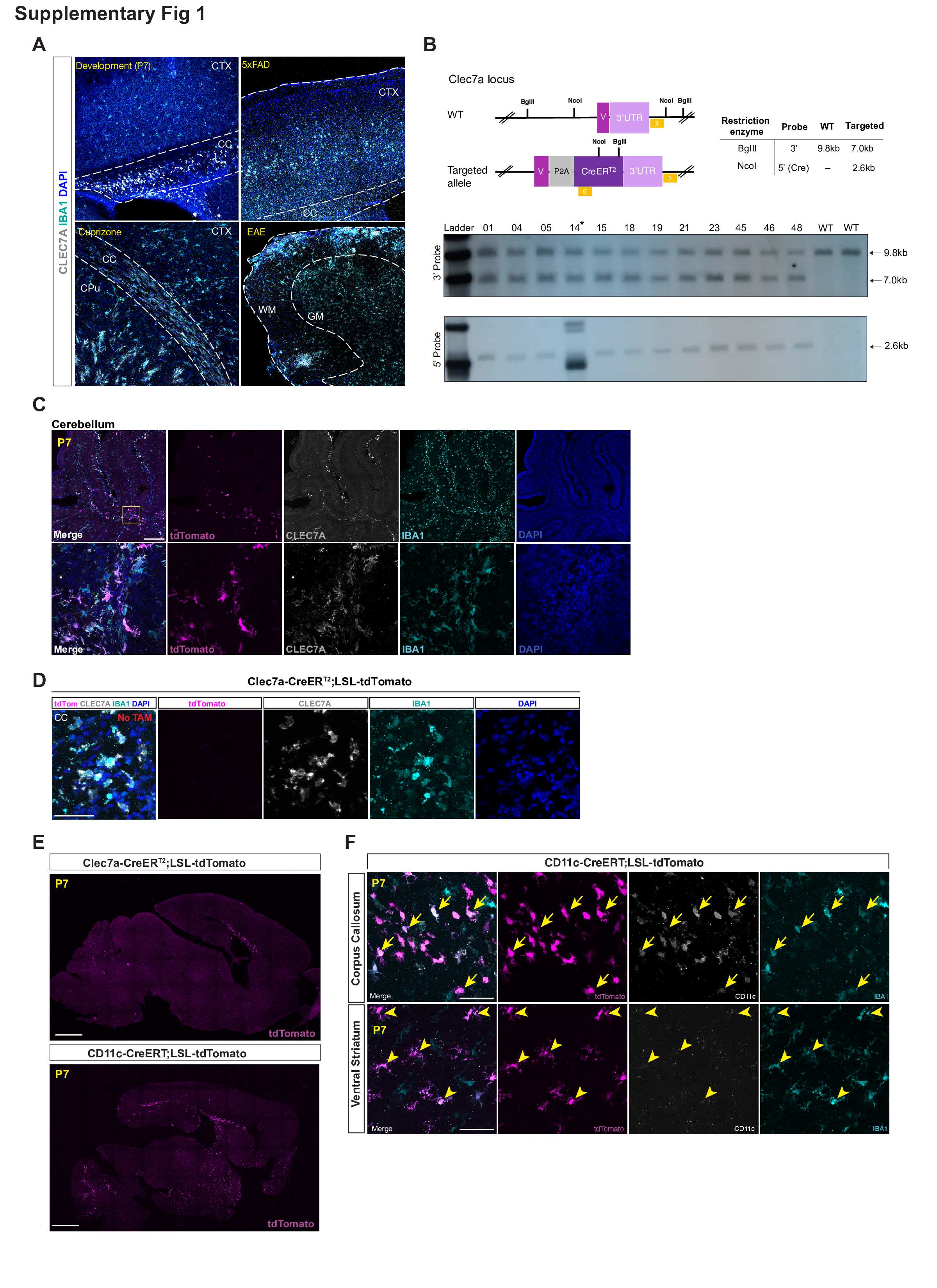

**Figure S1. Generation of Clec7a-CreER^T2^ mouse**

(A) Immunostaining validation of CLEC7A expression, which overlaps with microglial marker IBA1 signals, in development (P7), 5xFAD, cuprizone and EAE models, scale bar = 200um. Composite images for Figure 1A. CC, corpus callosum; CTX, cortex; HIP, hippocampus; CPu, caudate putamen; WM, white matter; GM, gray matter.
(B) Southern blot confirmation of a single CreER insertion into the targeted genomic locus in F1 animals (offspring of founders crossed with wildtype mice). Successful integration of a single CreER insertion downstream of the *Clec7a* coding region displays a 7.0kb band when using BglII digestion and 3’ probe detection, and a 2.6kb band when using NcoI digestion followed by 5’ (Cre) probe detection. Alleles with no insertion (WT) has a 9.8kb band when using BglII digestion and 3’ probe detection, and no band for the 5’ probe. Mouse ID# are labeled on the gel, showing that all lines tested except #14 have correct blotting patterns, and these lines are heterozygous for the transgene as expected.
(C) Representative immunostaining images showing that the Clec7a-CreER^T2^ reporter mouse labels PAM in cerebellar white matter of a P7 brain. tdTomato^+^ cells are almost all CLEC7A^+^IBA1^+^. Bottom panel displays a zoomed-in image of the boxed area in the top panel. Scale bar = 250um (top) and 50um (bottom).
(D) Representative immunostaining images showing no tdTomato^+^ labeling in P7 Clec7a-CreERT2;LSL-tdTomato mice without tamoxifen injection. Scale bar = 50um. CC, corpus callosum.
(E) Representative immunostaining images showing the Clec7a-CreER^T2^ reporter mouse is more specific than the existing CD11c-CreERT line for PAM labeling. Scale bar = 1000um.
(F) Representative immunostaining images showing that CD11c-CreERT;LSL-tdTomato not only labels PAM (IBA1^+^CD11c^+^) in the corpus callosum (arrows) but also labels microglia that are stained negative for CD11c in the ventral striatum (arrowheads) at P7. Scale bar = 50um.

**Figure S2**

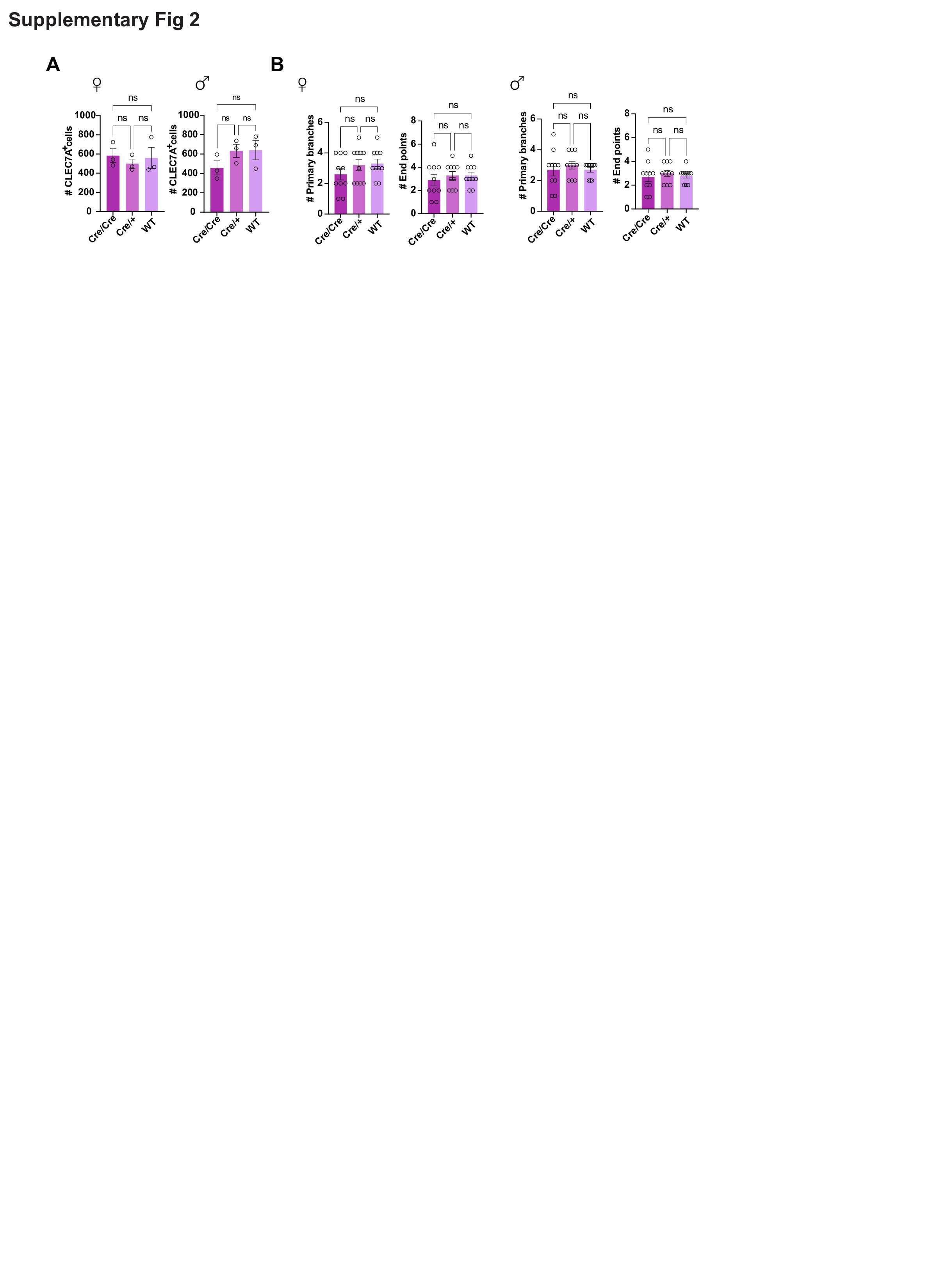
**Figure S2. Clec7a-CreER^T2^ does not affect *Clec7a* gene function**

(A) Quantification of Clec7a^+^ cells across Clec7a-CreER^T2^ homozygous, heterozygous and wildtype littermates split by sex. n = 3 mice (3 sections per animal). One-way ANOVA with Tukey’s multiple comparisons test. ns: not significant. Error bars represent mean +/- SEM.
(B) Quantification of primary branches (left) and end points (right) per cell across Clec7a-CreER^T2^ homozygous, heterozygous and wildtype littermates split by sex. n = 10 per group. One-way ANOVA with Tukey’s multiple comparisons test. ns: not significant. Error bars represent mean +/- SEM.

**Figure S3**
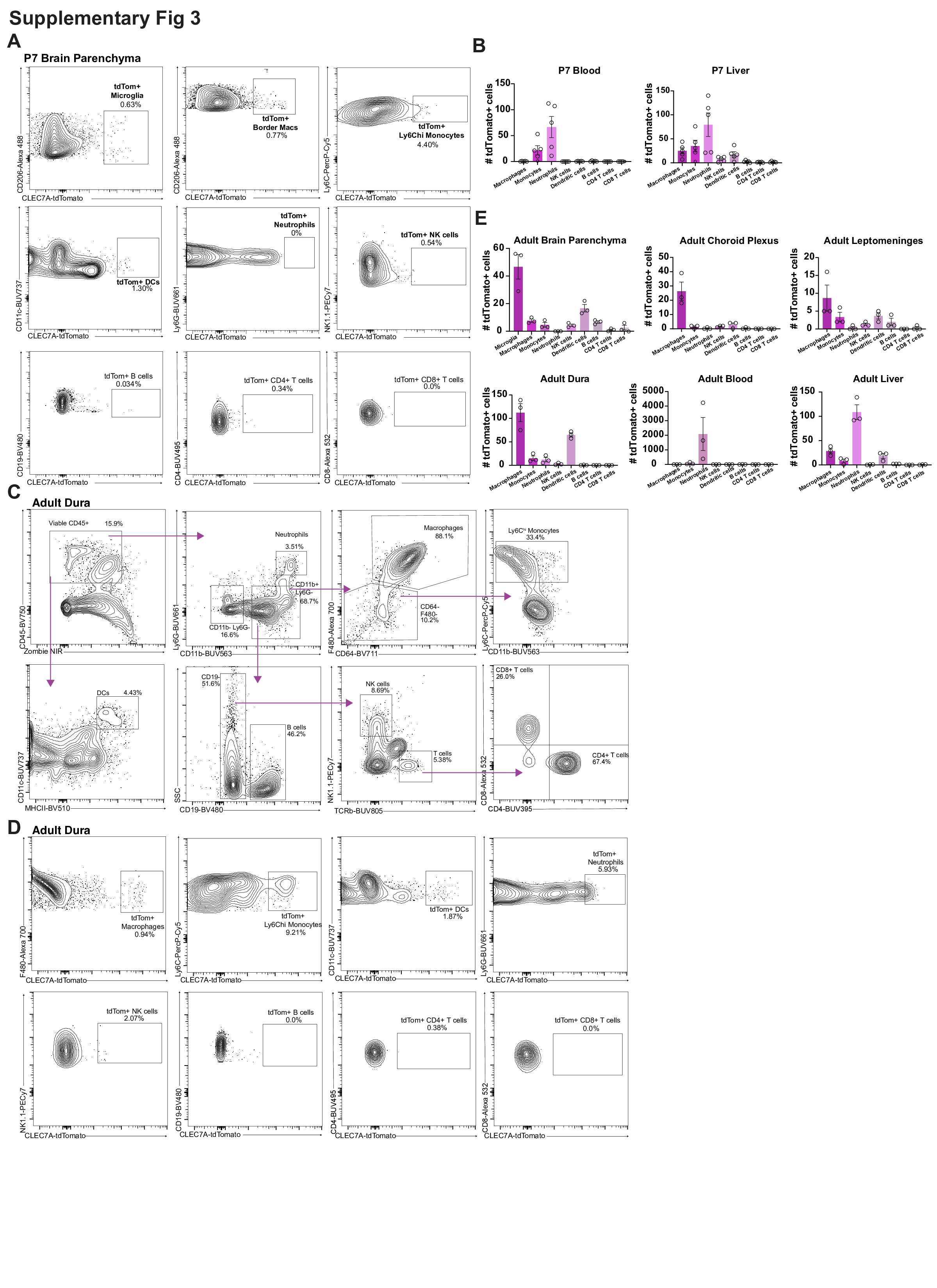

**Figure S3. Characterization of tdTomato^+^ immune cells in the brain and peripheral tissues in P7 and adult mice by flow cytometry.**

(A) Flow cytometry gating and quantification of immune cells expressing tdTomato reporter in P7 tissues.
(B) Quantification of tdTomato^+^ immune cells in P7 blood and liver. n = 5 mice. Error bars represent mean +/- SEM.
(C) Flow cytometry gating strategy for different immune cells in the Clec7a-CreER^T2^;LSL-tdTomato adult mice (represented by adult dura).
(D) Flow cytometry gating and quantification of immune cells expressing tdTomato reporter in adult tissues (represented by adult dura).
(E) Quantification of tdTomato^+^ immune cells in adult tissues. n = 3 mice. Error bars represent mean +/- SEM.

**Figure S4**
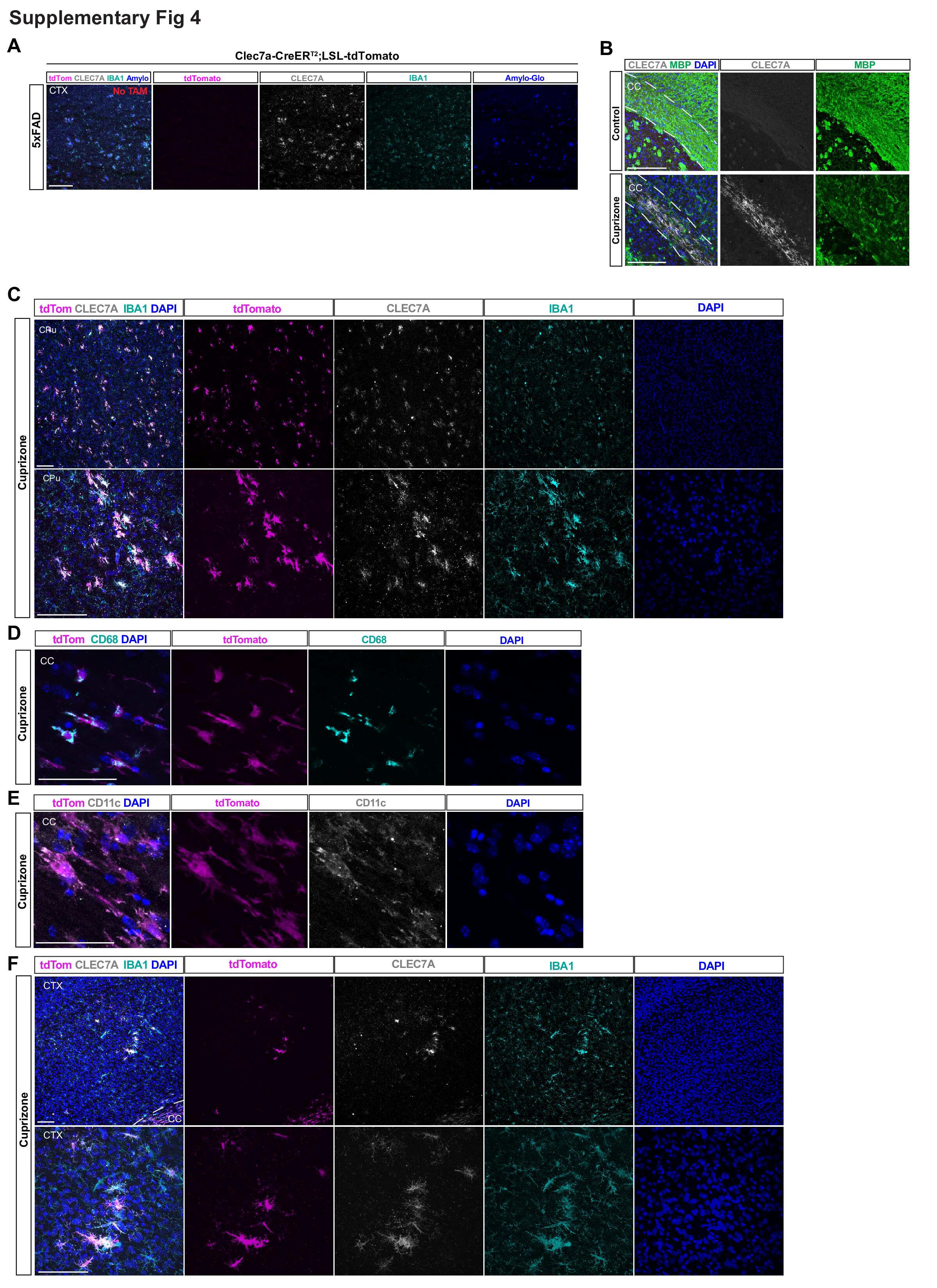

**Figure S4. Clec7a-CreER^T2^ reporter labels DAM across disease models**

(A) Representative immunostaining images showing no tdTomato^+^ labeling in 6-month Clec7a-CreER^T2^;LSL-tdTomato;5xFAD mice without tamoxifen injection. Scale bar = 50um. Scale bar = 200um. CTX, cortex.
(B) Representative images showing loss of myeline (MBP^+^) and appearance of CLEC7A^+^ microglia in corpus callosum upon cuprizone treatment (bottom). Scale bar = 200um.
(C) Representative images showing tdTomato^+^ microglia labeling in caudate putamen (CPu) of Clec7a-CreER^T2^;LSL-tdTomato mice treated with cuprizone. Scale bar = 200um (top, lower magnification) and 100um (bottom, higher magnification).
(D) Representative images showing tdTomato^+^ microglia co-labeled with CD68 in corpus callosum of Clec7a-CreER^T2^;LSL-tdTomato mice treated with cuprizone, Scale bar = 50um.
(E) Representative images showing tdTomato^+^ microglia co-labeled with CD11c in corpus callosum of Clec7a-CreER^T2^;LSL-tdTomato mice treated with cuprizone, Scale bar = 50um.
(F) Representative images showing rare tdTomato^+^ microglia in cortex (CTX) of Clec7a-CreER^T2^;LSL-tdTomato mice treated with cuprizone. Scale bar = 200um (top, lower magnification) and 100um (bottom, higher magnification).

**Figure S5**

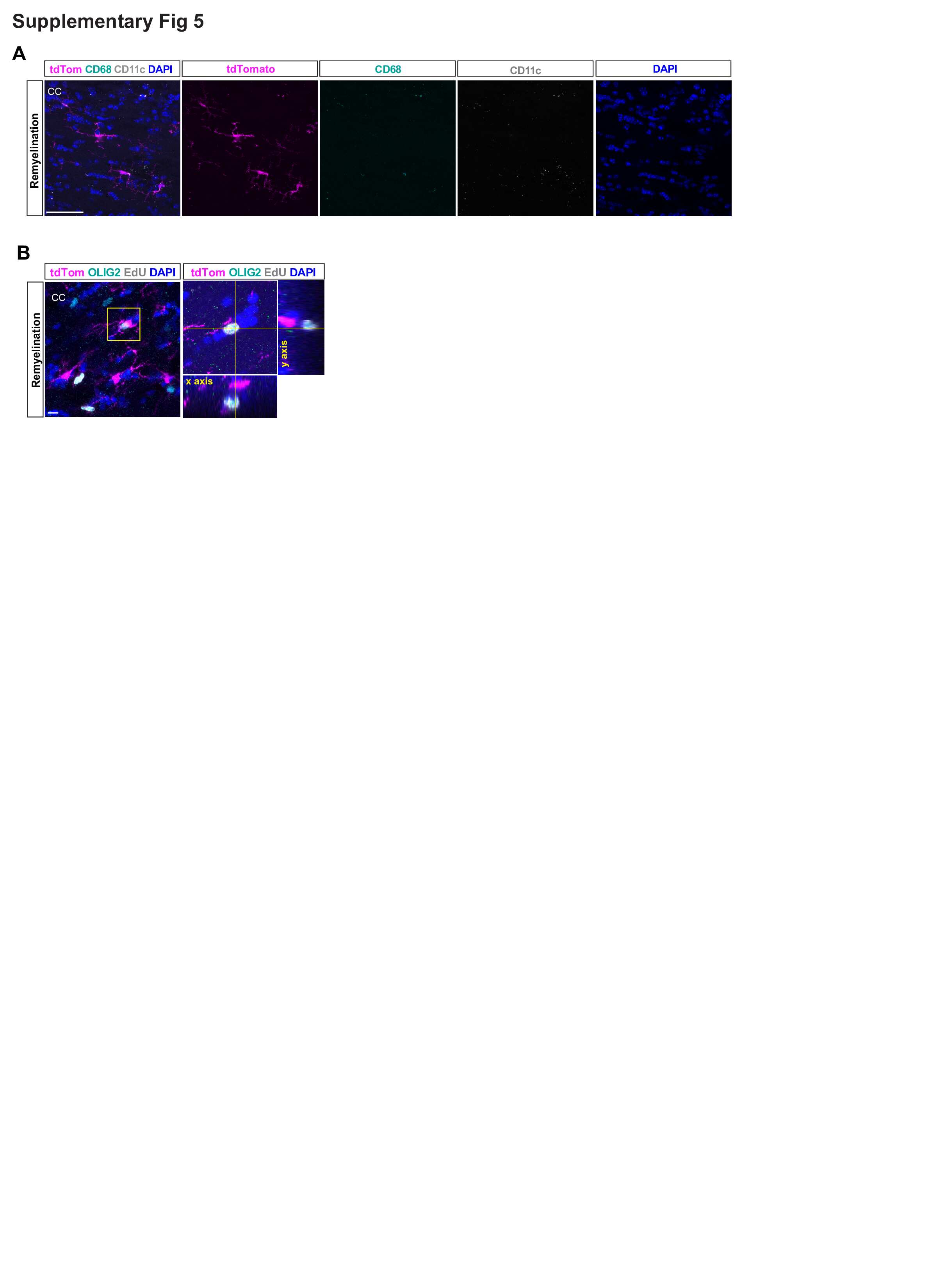

**Figure S5. DAM return to homeostasis during remyelination in cuprizone model**

(A) Representative images showing tdTomato^+^ cells downregulate CD68 and CD11c in corpus callosum following remyelination. Scale bar = 50um.
(B) X and Y orthogonal view of the boxed cell validating that the tdTomato^+^ cell does not overlap with EdU. Scale bar = 50um.

**Figure S6**
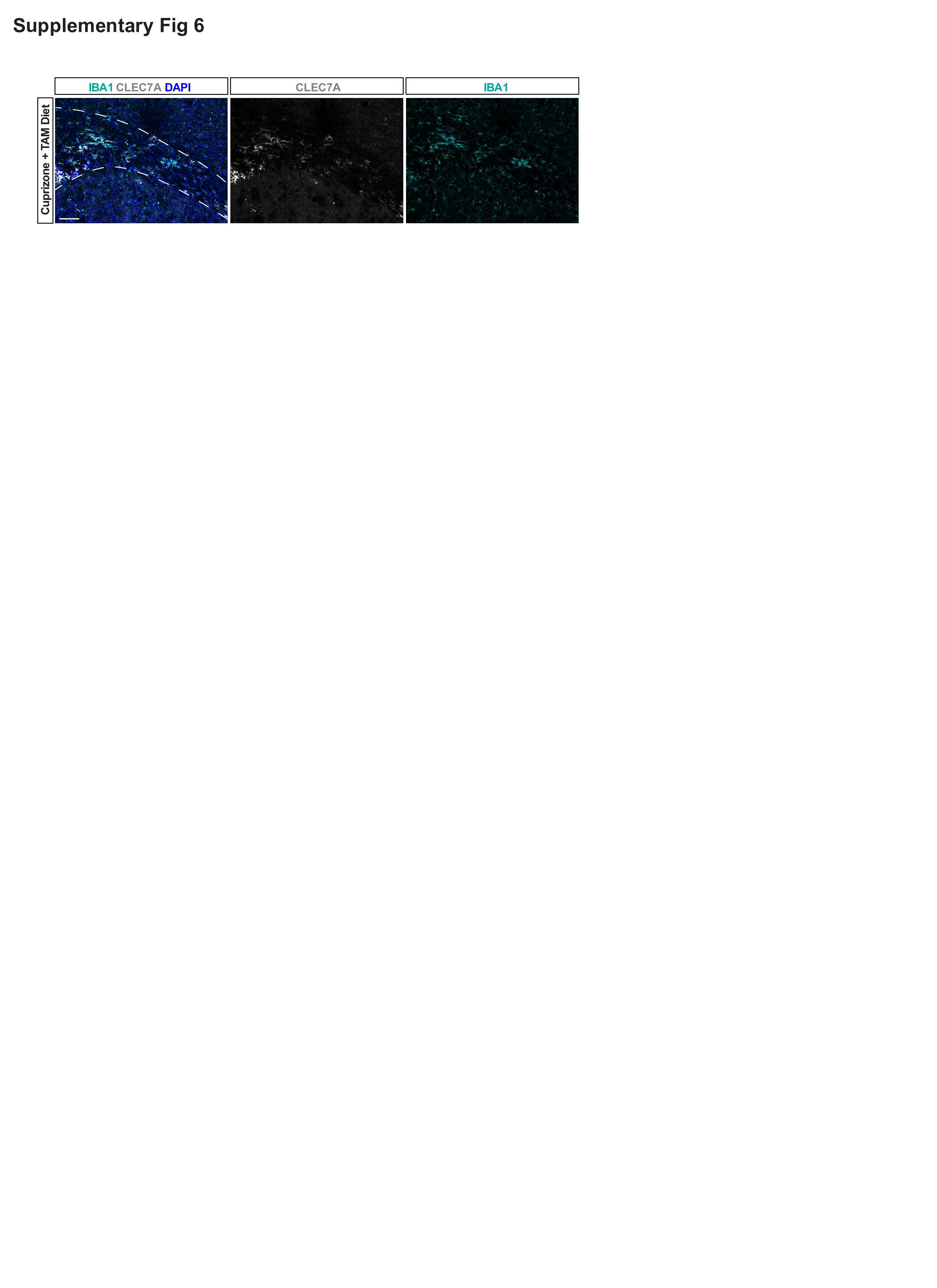

**Figure S6. DAM are present in the corpus callosum of control mice on cuprizone and tamoxifen diet.**

Representative images showing abundant CLEC7A^+^ microglia in the corpus callosum (CC) of Clec7a-creER^T2^;LSL-tdTomato mice treated with cuprizone and tamoxifen combined diet. Scale bar = 100um.
